# A recurrent network model of planning explains hippocampal replay and human behavior

**DOI:** 10.1101/2023.01.16.523429

**Authors:** Kristopher T. Jensen, Guillaume Hennequin, Marcelo G. Mattar

## Abstract

When faced with a novel situation, humans often spend substantial periods of time contemplating possible futures. For such planning to be rational, the benefits to behavior must compensate for the time spent thinking. Here we capture these features of human behavior by developing a neural network model where planning itself is controlled by prefrontal cortex. This model consists of a meta-reinforcement learning agent augmented with the ability to plan by sampling imagined action sequences from its own policy, which we call ‘rollouts’. The agent learns to plan when planning is beneficial, explaining empirical variability in human thinking times. Additionally, the patterns of policy rollouts employed by the artificial agent closely resemble patterns of rodent hippocampal replays recently recorded during spatial navigation. Our work provides a new theory of how the brain could implement planning through prefrontal-hippocampal interactions, where hippocampal replays are triggered by – and adaptively affect – prefrontal dynamics.

## Introduction

Humans and many other animals can adapt rapidly to new information and changing environments. Such adaptation often involves spending extended and variable periods of time contemplating possible futures before taking an action (Callaway et al., 2022; van Opheusden et al., 2023). For example, we might take a moment to think about which route to take to work depending on traffic conditions. The next day, some roads might be blocked due to roadworks, requiring us to adapt and mentally review the available routes in a process of re-planning before leaving the house. Since thinking does not involve the acquisition of new information or interactions with the environment, it is perhaps surprising that it is so ubiquitous for human decision making. However, thinking allows us to perform more computations with the available information, which can improve performance on downstream tasks (Bansal et al., 2022). Since physically interacting with the environment can consume time and other resources, or incur unnecessary risk, the benefits of planning often more than make up for the time spent on the planning process itself.

Despite a wealth of cognitive science research on the algorithmic underpinnings of planning (Solway and Botvinick, 2012; Callaway et al., 2022; Mattar and Daw, 2018; Mattar and Lengyel, 2022), little is known about the underlying neural mechanisms. This question has been difficult to address due to a scarcity of intracortical recordings during planning, and during contextual adaptation more generally. However, neuroscientists have begun to collect large-scale neural recordings during increasingly complex behaviors from the hippocampus and prefrontal cortex, brain regions known to be important for memory, decision making, and adaptation (Widloski and Foster, 2022; Pfeiffer and Foster, 2013; Gillespie et al., 2021; Wang et al., 2018; Samborska et al., 2022; Jadhav et al., 2016; Wu et al., 2017). These studies have demonstrated the importance of prefrontal cortex for generalizing abstract task structure across contexts (Wang et al., 2018; Samborska et al., 2022). Additionally, it has been suggested that planning could be mediated by the process of hippocampal forward replays (Pfeiffer and Foster, 2013; Widloski and Foster, 2022; Mattar and Daw, 2018; Agrawal et al., 2022; Foster, 2017; Jiang et al., 2022; Johnson and Redish, 2007). Despite these preliminary theories, little is known about how hippocampal replays could be integrated within the dynamics of downstream circuits to implement planning-based decision making and facilitate adaptive behavior (Yu and Frank, 2015).

While prevailing theories of learning from replays generally rely on dopamine-mediated synaptic plasticity (Gomperts et al., 2015; Mattar and Daw, 2018; De Lavilléon et al., 2015), it is currently unclear whether this process could operate sufficiently fast to also inform online decision making. It has recently been suggested that some forms of fast adaptation could result from recurrent meta-reinforcement learning (meta-RL; Wang et al., 2018, 2016; Duan et al., 2016). Such meta-RL models posit that adaptation to new tasks can be directly implemented by the recurrent dynamics of the prefrontal network. The dynamics themselves are learned through gradual changes in synaptic weights, which are modified over many different environments and tasks in a slow process of reinforcement learning. Importantly, such recurrent neural network (RNN)-based agents are able to adapt rapidly to a new task or environment after training by integrating their experiences into the hidden state of the RNN, with no additional synaptic changes (Wang et al., 2018, 2016; Duan et al., 2016; Zintgraf et al., 2019; Alver and Precup, 2021). However, previous models are generally only capable of making *instantaneous* decisions and thus do not have the ability to improve their choices by ‘thinking’ prior to taking an action. Wang et al. (2018) explored the possibility of allowing multiple steps of network dynamics before making a decision, but this additional computation was also pre-determined by the experimenter and not adaptively modulated by the agent itself.

In this work, we propose a model that similarly combines slow synaptic learning with fast adaptation through recurrent dynamics in the prefrontal network. In contrast to previous work, however, this recurrent meta-learner can *choose* to momentarily forgo physical interactions with the environment and instead ‘think’ (Hamrick et al., 2017; Pascanu et al., 2017). This process of thinking is formalized as the simulation of sequences of imagined actions, sampled from the policy of the agent itself, which we refer to as ‘rollouts’ (Figure 1A). We introduce a flexible maze navigation task to study the relationship between the behavior of such RL agents and that of humans (Figure 1B). RL agents trained on this task learn to use rollouts to improve their policy and better generalize to previously unseen environments, and they selectively trigger rollouts in situations where humans also spend more time deliberating.

**Figure 1:**
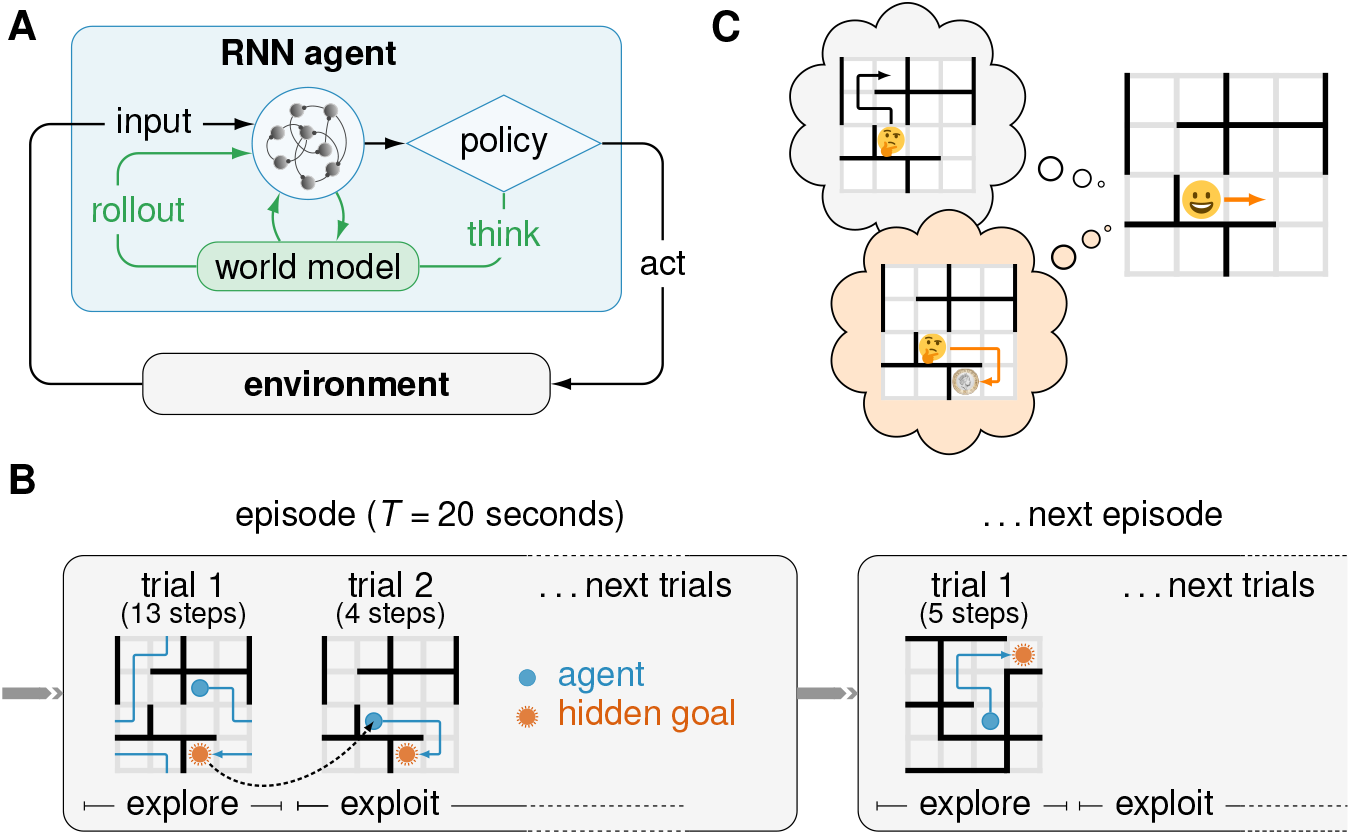
Task and model schematics. **(A)** The RL agent consisted of a recurrent neural network, which received information about the environment and executed actions in response. The primary output of the agent was a policy from which the next action was sampled. This action could either be to move in the environment in a given direction (up, down, left or right), or to ‘think’ by using an internal world model to simulate a possible future trajectory (a ‘rollout’). The agent was trained to maximize its average reward per episode and to predict (i) the upcoming state, (ii) the current goal location, and (iii) the value of the current state. When the agent decided to plan, the first two predictors were used in an open-loop planning process, where the agent iteratively sampled ‘imagined’ actions and predicted what the resulting state would be, and whether the goal had been (virtually) reached. The output of this planning process was appended to the agent’s input on the subsequent time step (details in text). A physical action was assumed to take 400 ms and a rollout was assumed to take 120 ms (Kurth-Nelson et al., 2016). **(B)** Schematic illustrating the dynamic maze task. In each episode lasting *T* = 20 seconds, a maze and a goal location were randomly sampled. Each time the goal was reached, the subject received a reward and was subsequently “teleported” to a new random location, from which it could return to the goal to receive more reward. The maze had periodic boundaries, meaning that subjects could exit one side of the maze to appear at the opposite side. **(C)** Schematic illustrating how policy rollouts can improve performance by altering the momentary policy. An agent might perform a policy rollout leading to low value (top; black), which would *decrease* the probability of physically performing the corresponding sequence of actions. Conversely, a rollout leading to high value (bottom; orange) would *increase* the probability of the corresponding action sequence. Notably, these policy changes occur at the level of network dynamics rather than parameter updates.

Additionally, we draw explicit parallels between the model rollouts and hippocampal replays through novel analyses of recent hippocampal recordings from rats performing a similar maze task (Widloski and Foster, 2022). The content and behavioral effects of hippocampal replays in this dataset have a striking resemblance to the content and effects of policy rollouts in our computational model. Our work thus addresses two key questions from previous studies on hippocampal replays and planning. First, we show that a recurrent network can meta-learn when to plan instead of having to precompute a ‘plan’ in order to decide whether to use it (Mattar and Daw, 2018; Russek et al., 2022). Second, we propose a new theory of replay-mediated planning, which utilizes fast network dynamics for real-time decision making that could operate in parallel to slower synaptic plasticity (Gomperts et al., 2015). To formalize this second point, we provide a normative mathematical theory of how replays can improve decision making via feedback to prefrontal cortex by approximating policy gradient optimization (Sutton and Barto, 2018). We show that such an optimization process naturally arises in our RL agent trained for rapid adaptation and suggest that biological replays could implement a similar process of rollout-driven decision making (Figure 1C).

Our work provides new insights into the neural underpinning of ‘thinking’ by bridging the gap between recurrent meta-RL (Wang et al., 2018), machine learning research on adaptive computation (Hamrick et al., 2017; Graves, 2016; Banino et al., 2021), and theories of meta-cognition (Griffiths et al., 2015; Botvinick and Cohen, 2014; Botvinick et al., 2020). We link these ideas to the phenomenon of hippocampal replay and provide a new theory of how forward replays can modulate behavior through recurrent interactions with prefrontal cortex.

## Results

### Humans think for different durations in different contexts

To characterize the behavioral signatures of planning, we recruited 94 human participants from Prolific to perform an online maze navigation task where the walls and goal location changed periodically. The environment was a 4 *×*4 grid with periodic boundaries, impassable walls, and a single hidden reward (Figure 1B; Methods; see Figure S1 for results with non-periodic boundaries). The task consisted of several ‘episodes’ lasting *T* = 20 seconds each. At the start of each episode, the wall configuration, reward location, and initial position were randomly sampled and fixed until the next episode. In the first trial, subjects explored the maze by taking discrete steps in the cardinal directions until finding the hidden reward. Subjects were then immediately moved to a new random location, initiating an exploitation phase where they had to repeatedly return to the same goal location from random start locations (Figure 1B). Participants were paid a monetary bonus proportional to the average number of trials completed per episode (Methods; Figure S2), and they displayed clear signs of learning in the form of increasing reward and decreasing response times over the 40 episodes of the experiment (Figure S3A-B).

We first examined human performance as a function of trial number within each episode, comparing the first exploration trial with subsequent exploitation trials. Participants exhibited a rapid ‘one-shot’ transition to goal-directed navigation after the initial exploration phase (Figure 2A, black), consistent with previous demonstrations of rapid adaptation in ‘meta-learning’ settings (Wang et al., 2018). We next investigated the time participants spent thinking during the exploitation phase. We estimated the ‘thinking time’ for each action as the posterior mean under a probabilistic model that decomposes the total response time for each action (Figure 2B; top) into the sum of the thinking time (Figure 2B; bottom) and a perception-action delay. The prior distribution over perception-action delays was estimated for each individual using a separate set of episodes, where participants were explicitly cued with the optimal path to eliminate the need for route planning (Methods; Figure S2). Since the first action within each trial also required participants to parse their new position in the maze, a separate prior was fitted for these actions.

**Figure 2:**
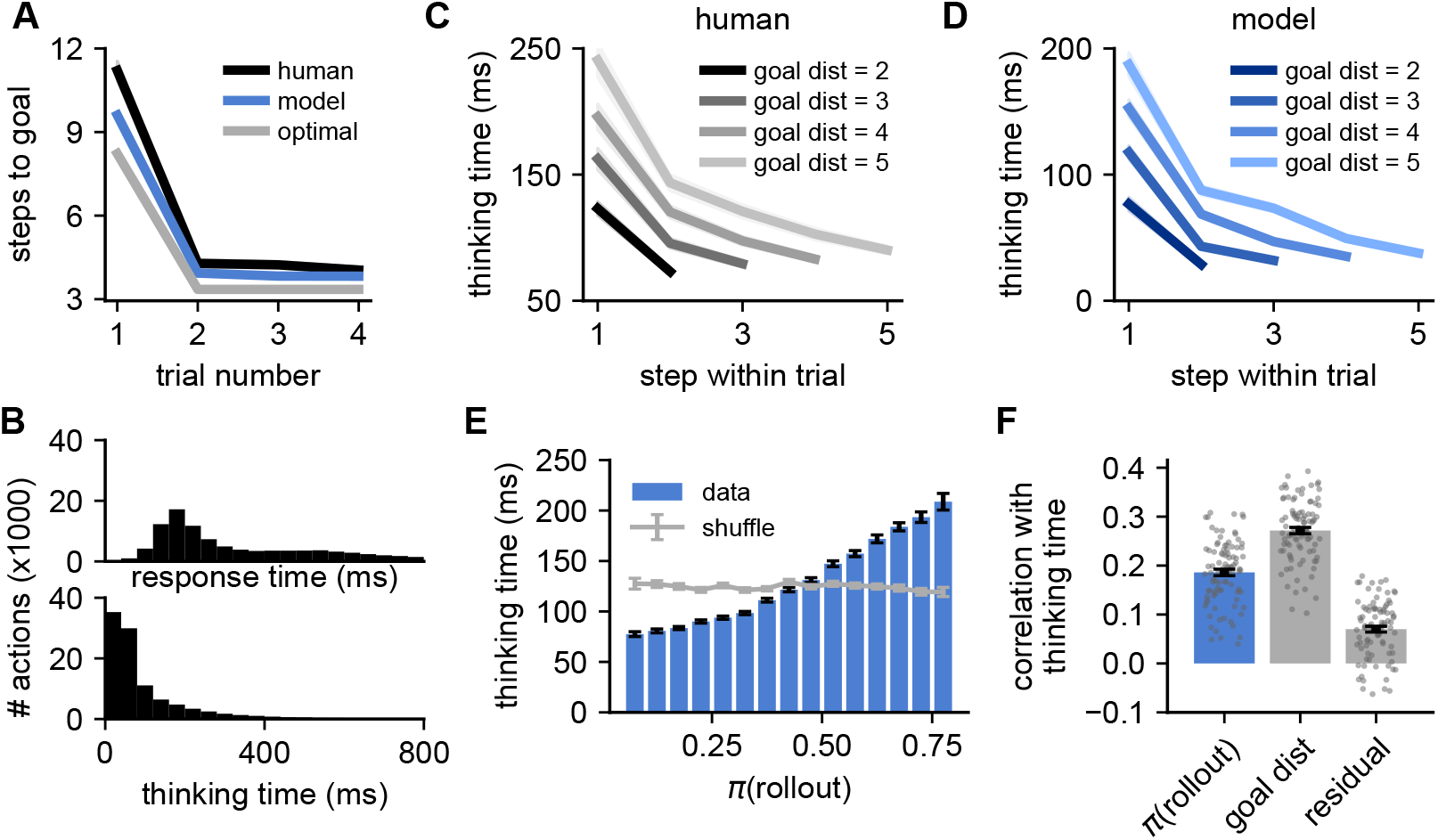
Trained RL agents perform more rollouts in situations where humans spend longer thinking. **(A)** Performance – quantified as the number of actions taken to reach the goal – as a function of trial number within each episode, computed for both human participants (black) and RL agents (blue). Shading indicates standard error of the mean across human participants (*n* = 94) or RL agents (*n* = 5) and mostly falls within the interval covered by the solid lines. Gray line indicates optimal performance, computed separately for exploration (trial 1) and exploitation (trials 2-4; Methods). **(B)** Distribution of human response times (top) and thinking times (bottom), spanning ranges on the order of a second (Methods). **(C)** Human thinking time as a function of the step-within-trial (x-axis) for different initial distances to the goal at the beginning of the trial (lines, legend). Shading indicates standard error of the mean across 94 participants. Participants spent more time thinking further from the goal and before the first action of each trial (Figure S4). **(D)** Model ‘thinking times’ separated by time-within-trial and initial distance-to-goal, exhibiting a similar pattern to human participants. To compute thinking times for the model, each rollout was assumed to last 120 ms as described in the main text. Shading indicates standard error of the mean across five RL agents. The average thinking time can be less than 120 ms because the agents only perform rollouts in some instances and otherwise make a ‘reflexive’ decision. This is particularly frequent near the goal and late in a trial, where humans also spend less time thinking. **(E)** Binned human thinking time as a function of the probability that the agent chooses to perform a rollout, *π*(rollout). Error bars indicate standard error of the mean within each bin. Gray horizontal line indicates a shuffled control, where human thinking times were randomly permuted before the analysis. **(F)** Correlation between human thinking time and the regressors (i) *π*(rollout) under the model, (ii) momentary distance-to-goal, and (iii) *π*(rollout) after conditioning on the momentary distance-to-goal (‘residual’; Methods). Bars and error bars indicate mean and standard error across human participants, and gray dots indicate individual participants (*n* = 94).

Participants exhibited a wide distribution of thinking times during the exploitation phase of the task (Figure 2B; bottom). To reveal any task-related structure in this variability, we partitioned thinking times by within-trial action number and by the initial distance-to-goal for the trial (Figure 2C). Thinking times were longer when further from the goal, consistent with planning of longer routes taking more time. Thinking times were also substantially longer for the first action of each trial (Figure S4), consistent with participants having to initially plan a new route to the goal. These patterns confirm that the broad marginal distribution of thinking times (Figure 2B) does not simply reflect a noisy decision making process or task-irrelevant distractions. On the contrary, variability in thinking time is an important feature of human behavior that reflects the variable moment-to-moment cognitive demands for decision making.

### A recurrent network model of planning

To model the rapid adaptation and the detailed patterns of thinking times displayed by human subjects, we considered an RNN model trained in a meta-reinforcement learning setting (Figure 1A; Methods; Duan et al., 2016; Wang et al., 2016, 2018; see Supplementary Note for a more in-depth discussion and motivation of various modeling choices). The RL agent had 100 gated recurrent units (GRUs; Cho et al., 2014; Figure S5), whose time-varying internal state ***h***_*k*_ evolved dynamically according to

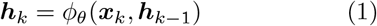

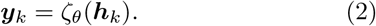

*θ* denotes the model parameters, ***x***_*k*_ are RNN inputs, and ***y***_*k*_ are its outputs. ***h***_*k*_ was reset at the beginning of each episode. *k* indexes the evolution of the network dynamics, which can differ from the wall-clock time *t* in agents augmented with the ability to ‘think’ (see below). Inputs consisted of the current agent location *s*_*k*_, previous action *a*_*k−*1_, reward *r*_*k−*1_, wall locations, and the elapsed time *t* since the start of the episode (Methods). While the reward location was hidden and had to be discovered, the rest of the environment was fully observed. The output consisted primarily of a policy *π*_*θ*_(*a*_*k*_ | ***h***_*k*_), which was a function of the network state. At each iteration, an action *a*_*k*_ was sampled from *π*_*θ*_(*a*_*k*_ | ***h***_*k*_). This triggered environment changes ***x***_*k*+1_, *s*_*k*+1_ = *ψ*(*a*_*k*_, *s*_*k*_), which resulted in a new location *s*_*k*+1_ and inputs ***x***_*k*+1_ that were fed back to the agent (Figure 1A). In addition to the policy, the RNN output included a value function (Figure S6) and predictions of the agent’s next location and the current goal location (Figure S7).

Performance was quantified as the expected total reward,

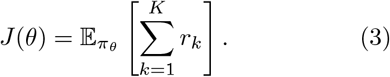

*K* denotes the number of iterations per episode, with each episode terminating when *t* exceeded *T* = 20 seconds as in the human data (Figure 1B). During training, the parameters *θ* were adjusted using policy gradients to maximize the average *J* (*θ*) across environments (Methods; Sutton and Barto, 2018; Wang et al., 2018; Jensen, 2023b). Since the agent lacked an intrinsic notion of wallclock time, we considered each action to consume Δ*t* = 400 ms. This allowed 50 actions per episode, similar to the human data (Supplementary Note).

In this canonical formulation, the agent always takes an instantaneous action in response to its inputs, implying constant (zero) ‘thinking time’ in all situations. This formulation can therefore not explain the salient patterns of thinking times observed in human participants (recall Figure 2C). Temporally extended planning might also appear *unnecessary*, since the agent has access to all information required for decision making, including the current state, wall configuration, and reward location. However, this was also true for human participants, who spent time thinking nonetheless. We hypothesized that the RL agent could similarly benefit from the ability to trade off time for additional *processing* of the available information (Hamrick et al., 2017; Pascanu et al., 2017; Supplementary Note).

To test this hypothesis, we augmented the RL agent with the ability to perform temporally extended planning in the form of imagined policy rollouts. Specifically, we expanded the action space of the agent to give it the option of sampling a hypothetical trajectory from its own policy at any moment in time (a ‘rollout’; Figure 1A; Hamrick et al., 2017; Pascanu et al., 2017). In other words, the agent was allowed to either perform a physical action, or to perform a mental simulation of its policy. A rollout took the form of a sequence of separate recurrent processing steps. At each step, the network ‘imagined’ taking an action sampled from its momentary policy at that step, and it used a learned world model (Figure 1A; see below) to predict the consequences of that action. In particular, the world model predicted the input that the RNN would normally receive at the next step if the imagined action were actually carried out from the imagined state. This predicted input was then used for the next step of recurrent processing in the rollout. The rollout process stopped after 8 imagined actions, or earlier if the agent imagined having hit the goal during the rollout (Supplementary Note; see Figure S5 for different network sizes and planning horizons).

To capture the fact that mental simulation is faster than physical actions (Liu et al., 2019; Kurth-Nelson et al., 2016), we assumed each full rollout of up to 8 imagined actions to consume only 120 ms (see Figure S8 for an alternative model where the temporal cost of a rollout is proportional to its length). In other words, a single iteration of the network dynamics (*k →k* + 1 in Equation 1) incremented time by 120 ms if the agent chose to perform a rollout and 400 ms if the agent chose a physical action. This allowed the agent to perform many simulated actions in the time it would take to physically move only a short distance (Agrawal et al., 2022). Importantly, since an episode had a fixed duration of 20 seconds, choosing to perform more rollouts had a temporal opportunity cost by leaving less time for physical actions towards the goal.

If the agent chose to perform a rollout, a flattened array of the imagined action sequence was fed back to the network as additional inputs on the subsequent iteration, together with an indication of whether or not the simulated action sequence reached the goal (Supplementary Note). These inputs in turn affected the policy by modulating ***h***_*k*_ through learnable input weights (Figure 1A). This is reminiscent of canonical RL algorithms that change their *parameters θ* to yield a new and improved policy on the basis of trajectories sampled from the current policy.

In our formulation of planning, the agent’s policy is instead induced by the *hidden state* ***h***_*k*_, which can similarly be modulated based on the imagined policy rollouts to improve performance.

Importantly, both the generation of a mental roll-out and the corresponding success feedback relied on an *internal* model of the environment that was obtained from the agent itself. This internal model was trained alongside the RNN and the policy by learning to predict the reward location and state transitions from the momentary hidden state of the RNN (***h***_*k*_) and the action taken (*a*_*k*_; Methods; Figure S7). At the beginning of each rollout, the most likely goal location according to the internal model was identified and used as an ‘imagined goal’ throughout the rollout. Rollouts thus did not provide any privileged information that the agent did not already possess. Instead, they allowed the agent to trade off time for additional computation – similar to thinking in humans and other animals.

Biologically, we interpret rollouts as prefrontal cortex (the RNN) interacting with the hippocampal formation (the world model) to simulate and evaluate an action sequence via replay. Following Wang et al. (2018), we use ‘prefrontal cortex’ as a general term for both PFC itself and associated areas of striatum and thalamus (Supplementary Note). Importantly, while we endowed the agent with the ability to perform policy rollouts, we did not build in any knowledge of when, how, or how much to use them. The agent instead learned this through training on many different environments. Therefore, while rollouts phenomenologically resembled hippocampal forward replays *by design*, our model allowed us to investigate (i) whether and how rollouts can drive policy improvements, (ii) whether their temporal patterns explain human response times, and (iii) whether biological replays might implement a similar computation.

The RL agent was trained by adjusting its parameters *θ* over 8 *×*10^6^ episodes, sampled randomly from 2.7 *×*10^8^ possible environment configurations. This large task space required the agent to generalize across tasks. Parameter changes followed the gradient of a cost function designed to (i) maximize expected reward (Equation 3), (ii) learn the internal model by predicting the reward location and state transitions, and (iii) maximize the policy entropy to encourage exploration (Methods; Wang et al., 2016). Importantly, parameters were frozen after training, and the agent adapted to new environments using only its network dynamics (Wang et al., 2018).

### Human thinking times correlate with agent rollouts

Having specified our computational model of planning, we analyzed its behavior and compared it to that exhibited by humans. We trained five copies of our RL agent to solve the same task as the human participants and found that the agents robustly learned to navigate the changing maze (Figure S3C). Similar to humans, the trained agents exhibited a rapid transition from exploration to exploitation upon finding the reward, reaching near-optimal performance in both phases (Figure 2A, blue). This confirmed that these RNNs are capable of adapting to changing environments using only internal network dynamics with fixed parameters, corroborating previous work on recurrent meta-RL (Wang et al., 2018; Duan et al., 2016; Banino et al., 2018). However, while the RL agents learned this structure through repeated exposure to the task, humans were immediately able to solve the task on the basis of written instructions (Figure S3A) – a potentially different type of ‘meta-learning’.

The trained networks also used their capacity to perform rollouts, choosing to do so on approximately 30% of all iterations after training (Figure S3D). Importantly, there was temporal variability in the probability of performing a rollout, and the networks sometimes performed multiple successive rollouts between consecutive physical actions. When we queried the conditions under which the trained agents performed these rollouts, we found striking similarities with the pattern of human thinking times observed previously. In particular, the RL agent performed more rollouts earlier in a trial and further from the goal (Figure 2D) – situations where the human participants also spent more time thinking before taking an action (Figure 2C). On average, thinking times in the RL agent were approximately 50 ms lower than in humans. This difference could e.g. be due to (i) differences in how the periodic boundaries are represented in humans and RL agents (Jensen, 2023a), (ii) the agent having a better ‘base policy’ than humans, or (iii) the hyperparameters determining the temporal cost of planning (Supplementary Note).

To further study the relationship between rollouts and human ‘thinking’, we simulated the RL agent in the same environments as the human participants. We did this by clamping the physical actions of the agent to those taken by the participants, while still allowing it to sample on-policy rollouts (Methods). In this setting, the agent’s probability of choosing to perform a rollout when encountering a new state, *π*(rollout), was a monotonically increasing function of human thinking time in the same situation (Figure 2E). The Pearson correlation between these two quantities was *r* = 0.186 *±* 0.007 (mean *±*sem across participants), which was significantly higher than expected by chance (Figure 2F, ‘*π*(rollout)’; chance level *r* = 0 *±*0.004). An above-chance correlation between thinking times and *π*(rollout) of *r* = 0.070 *±*0.006 persisted after conditioning on the momentary distance-to-goal (Figure 2F, ‘residual’), which was also correlated with thinking times (*r* = 0.272 *±*0.006). The similarity between planning in humans and RL agents thus extends beyond this salient feature of the task, including an increased tendency to plan on the first step of a trial (Figure S4).

In addition to the similarities during the exploitation phase, a significant correlation was also observed between human thinking time and *π*(rollout) during exploration (*r* = 0.098 *±*0.008). In this phase, both humans and RL agents spent more time thinking during later stages of exploration (Fig-ure S9). Model rollouts during exploration corresponded to planning towards an imagined goal from the posterior over goal locations, which becomes narrower as more states are explored (Figure S9). This finding suggests that humans may similarly engage in increasingly goal-directed behavior as the goal posterior becomes narrower over the course of exploration. In summary, a meta-reinforcement learning agent, endowed with the ability to perform rollouts, learns to do so in situations similar to when humans appear to plan. This provides a putative normative explanation for the variability in human thinking times observed in the dynamic maze task.

### Rollouts improve the policy of the RL agent

In the previous section, we saw that an RL agent can learn to use policy rollouts as part of its decision making process, and that the timing and number of rollouts correlate with variability in human thinking times. In this section, we aim to understand *why* the agent chooses to perform rollouts and *how* they guide its behavior. To do this, we considered the agent right after it first located the goal in each episode (i.e., at the first time step of trial 2; Figure 1B) and forced it to perform a pre-defined number of rollouts, which we varied. We then counted the number of actions needed to return to the goal while preventing any further rollouts during this return phase (Methods).

The average number of actions needed to reach the goal decreased monotonically as the number of forced rollouts increased up to at least 15 rollouts (Figure 3A). Interestingly, this was the case despite the unperturbed behavior of the agent rarely including more than a few consecutive rollouts (Figure 2D), suggesting that the agent learned a robust algorithm for policy optimization on the basis of such rollouts (Schrittwieser et al., 2020; Hamrick et al., 2017). To confirm that this performance improvement depended on the information contained in the policy rollouts rather than being driven by additional iterations of recurrent network dynamics, we repeated the analysis with no feedback from the rollout to the RNN and found a much weaker effect (Figure 3A, dashed gray line). The increase in performance with rollout number was also associated with a concomitant decrease in policy entropy (Figure 3B). Thus, performing more rollouts both improved performance and reduced uncertainty (Methods). These findings confirm that the agent successfully learned to use policy rollouts to optimize its future behavior. However, the question remains of whether this policy improvement is appropriately balanced with the temporal opportunity cost of performing a rollout.

**Figure 3:**
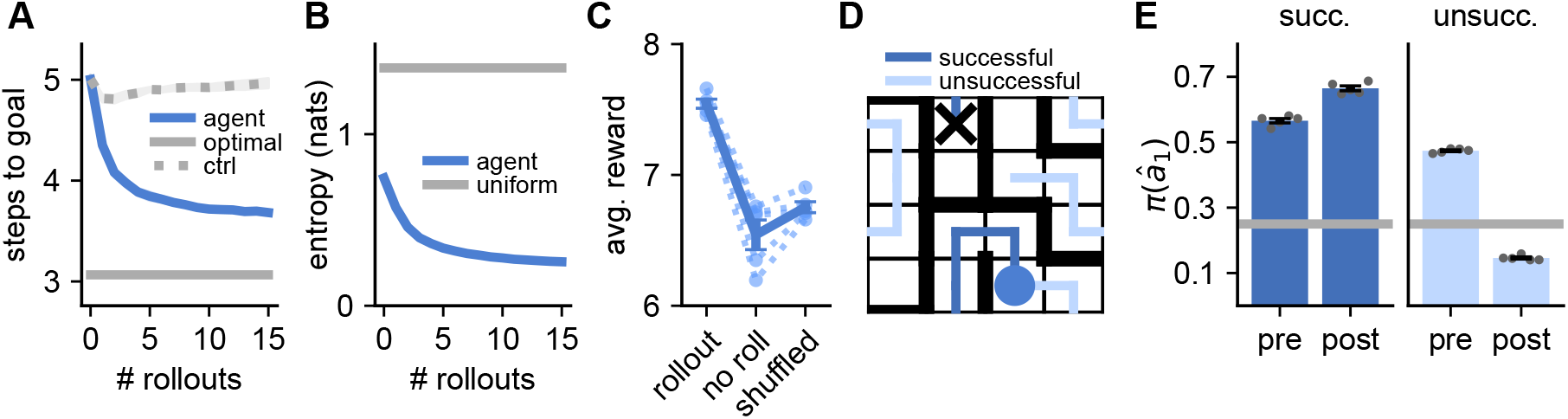
Rollouts improve the network policy. **(A)** Performance on trial 2 as a function of the number of rollouts enforced at the beginning of the trial, averaged over all trials. Performance was quantified as the average number of steps needed to reach the goal in the absence of further rollouts. Solid gray line (‘optimal’) indicates optimal performance, corresponding to the average initial distance-to-goal across trials. Dashed gray line (‘ctrl’) indicates a control simulation where the indicated number of rollouts was performed, but with the feedback to the RNN from the rollout channels set to zero. The performance gap from the non-perturbed agent confirms that the performance improvement with increasing numbers of rollouts is dependent on the information contained in the rollouts rather than being driven by additional iterations of recurrent network dynamics. **(B)** Policy entropy as a function of the number of rollouts enforced at the beginning of trial 2. The entropy was computed after re-normalizing the policy over the four ‘physical’ actions, and the horizontal gray line indicates the entropy of a uniform policy. **(C)** Original performance of the RL agent (left), performance when re-normalizing the policy over physical actions to prevent any rollouts (center), and performance after shuffling the timing of the rollouts while keeping the number of rollouts constant (right). Performance was quantified as the average number of rewards collected per episode, and dashed lines indicate individual RL agents, while the solid line indicates mean and standard error across agents. **(D)** Schematic showing an example of a ‘successful’ (dark blue) and an ‘unsuccessful’ (light blue) rollout from the same physical location (blue circle). Black cross indicates the goal location (not visible to the agent or human participants). **(E)** Probability of taking the first simulated action of the rollout, *â*_1_, before (*π*^pre^(*â*_1_)) and after (*π*^post^(*â*_1_)) the rollout. This was evaluated separately for successful (left) and unsuccessful (right) rollouts. *π*^pre^(*â*_1_) was above chance (gray line) in both cases and increased for successful rollouts, while it decreased for unsuccessful rollouts. Bars and error bars represent mean and standard error across five RL agents (gray dots). The magnitude of the changes in *π*(*â*_1_) depended on the planning horizon (Figure S5).

In general, rollouts are beneficial in situations where the policy improvement resulting from a rollout is greater than the temporal cost of 120 ms of performing the rollout. To investigate whether the agent learned to trade off the cost and benefits of rollouts (Hamrick et al., 2017; Pascanu et al., 2017; Agrawal et al., 2022), we computed the performance of the agent while explicitly forbidding rollouts. In this setting, each action was instead sampled from the distribution over physical actions only (Methods). When preventing rollouts in this way, the agent only collected 6.54*±* 0.11 rewards per episode compared to 7.54 *±*0.03 in the presence of rollouts (mean *±*standard error across agents), confirming that it used rollouts to increase expected reward (Figure 3C). To investigate whether the temporal structure of rollouts described in Figure 2 was important for this performance improvement, we performed an additional control, where the number of rollouts was kept fixed for each environment, but their occurrence was randomized in time. In this case, performance dropped to 6.75 *±*0.04 rewards per episode (Figure 3C), confirming that the RL agent used rollouts specifically when they improved performance.

To further dissect the effect of rollouts on agent behavior, we classified each rollout, 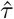 (a sequence *{â*_1_, *â*_2_, … *}* of rolled-out actions), as being either ‘successful’ if it reached the goal according to the agent’s internal world model, or ‘unsuccessful’ if it did not (Figure 3D). We hypothesized that the policy improvement observed in Figure 3A could arise from *upregulating* the probability of following a successful rollout and *downregulating* the probability of following an unsuccessful rollout. To test this hypothesis, we enforced a single rollout after the agent first found the reward and analyzed the effect of this rollout on the policy, separating the analysis by successful and unsuccessful rollouts. Importantly, we could compare the causal effect of rollout success by matching the history of the agent and performing rejection sampling from the rollout process until either a successful or an unsuccessful rollout had occurred (Methods). Specifically, we asked how the rollout affected the probability of taking the first rolled-out action, *â*_1_, by comparing the value of this probability before (*π*^pre^(*â*_1_)) and after (*π*^post^(*â*_1_)) the rollout. *π*^pre^(*â*_1_) was slightly higher for successful rollouts than unsuccessful rollouts, with both types of rollouts exhibiting a substantially higher-than-chance probability – a consequence of the model rollouts being drawn ‘on-policy’ (Figure 3E). However, while successful rollouts *increased π*(*â*_1_), unsuccessful rollouts *decreased π*(*â*_1_) (Figure 3E). This finding demonstrates that the agent combines the spatial information of a rollout with knowledge about its consequences, based on its internal world model, to guide future behavior.

### Hippocampal replays resemble policy rollouts

In our computational model, we designed policy rollouts to take the form of spatial trajectories that the agent could subsequently follow, and to occur only when the agent was stationary. These two properties are also important signatures of forward hippocampal replays – patterns of neural activity observed using electrophysiological recordings from rodents during spatial navigation (Pfeiffer and Foster, 2013; Widloski and Foster, 2022; Gillespie et al., 2021). Our model therefore allowed us to investigate whether forward replay in biological agents could serve a similar function during decision making to the function of policy rollouts in our RL agent. Additionally, since we have direct access to the agent’s policy and how it changes after a replay, our computational model can provide insights into the apparently conflicting data and contradictory viewpoints in the literature regarding the role of hippocampal replays. In particular, some studies have found a significant correlation between forward replay and subsequent behavior (Pfeiffer and Foster, 2013; Foster, 2017; Widloski and Foster, 2022), arguing that such a correlation suggests a role of forward replay for planning. On the contrary, other studies have found that forward replays do not always resemble subsequent behavior (Gillespie et al., 2021; Krause and Drugowitsch, 2022; Wu et al., 2017), challenging the interpretation of forward replay as a form of planning. Our model offers a potentially conciliatory explanation, predicting that the correlation between forward replay and subsequent behavior can be positive or negative, depending on the replayed trajectory (Figure 3E; Yu and Frank, 2015; Antonov et al., 2022).

To investigate whether there is evidence for such replay-based modulation of animal behavior, we reanalyzed a recently published hippocampal dataset from rats navigating a dynamic maze similar to the task in Figure 1B (Widloski and Foster, 2022). Animals had to repeatedly return to an initially unknown ‘home’ location, akin to the goal in our task (Figure S10). Both this home location and the configuration of the maze changed between sessions. Whilst the rats could not be ‘teleported’ between trials as in our task, they instead had to navigate to an unknown rewarded ‘away’ location selected at random after each ‘home’ trial. These ‘away’ trials served as a useful control since the animals did not know the location of the rewarded well at the beginning of the trial. Unlike the human data (Figure 2C), we found no correlation between the momentary distance-to-goal of the animal and time spent at the previously rewarded location (Figure S11). We hypothesize that this is because (i) the animals had to spend time consuming reward before they could continue, and (ii) a delay was imposed between reward consumption and the next reward becoming available. These periods could potentially be used for planning without incurring a substantial temporal opportunity cost, unlike the human task which explicitly enforced a trade-off between the time spent thinking and acting.

We therefore focused on the spatiotemporal content of hippocampal replays following previous hypotheses that they could form a neural substrate of planning (Pfeiffer and Foster, 2013; Foster, 2017; Widloski and Foster, 2022). We studied replay events detected in hippocampal recordings made with tetrode drives during the maze task (*n∈* [187, 333] simultaneously recorded neurons per session; Figure S10C). To detect replays, we followed Widloski and Foster (2022) and first trained a Bayesian decoder to estimate the animal’s position on a discretized grid from the neural data during epochs when the animal was moving. We then applied this decoder during epochs when the animal was stationary at a reward location before initiating a new trial, and defined replays as consecutive sequences of at least three adjacent decoded grid locations (Figure 4A; Figure S10; see Methods for details).

**Figure 4:**
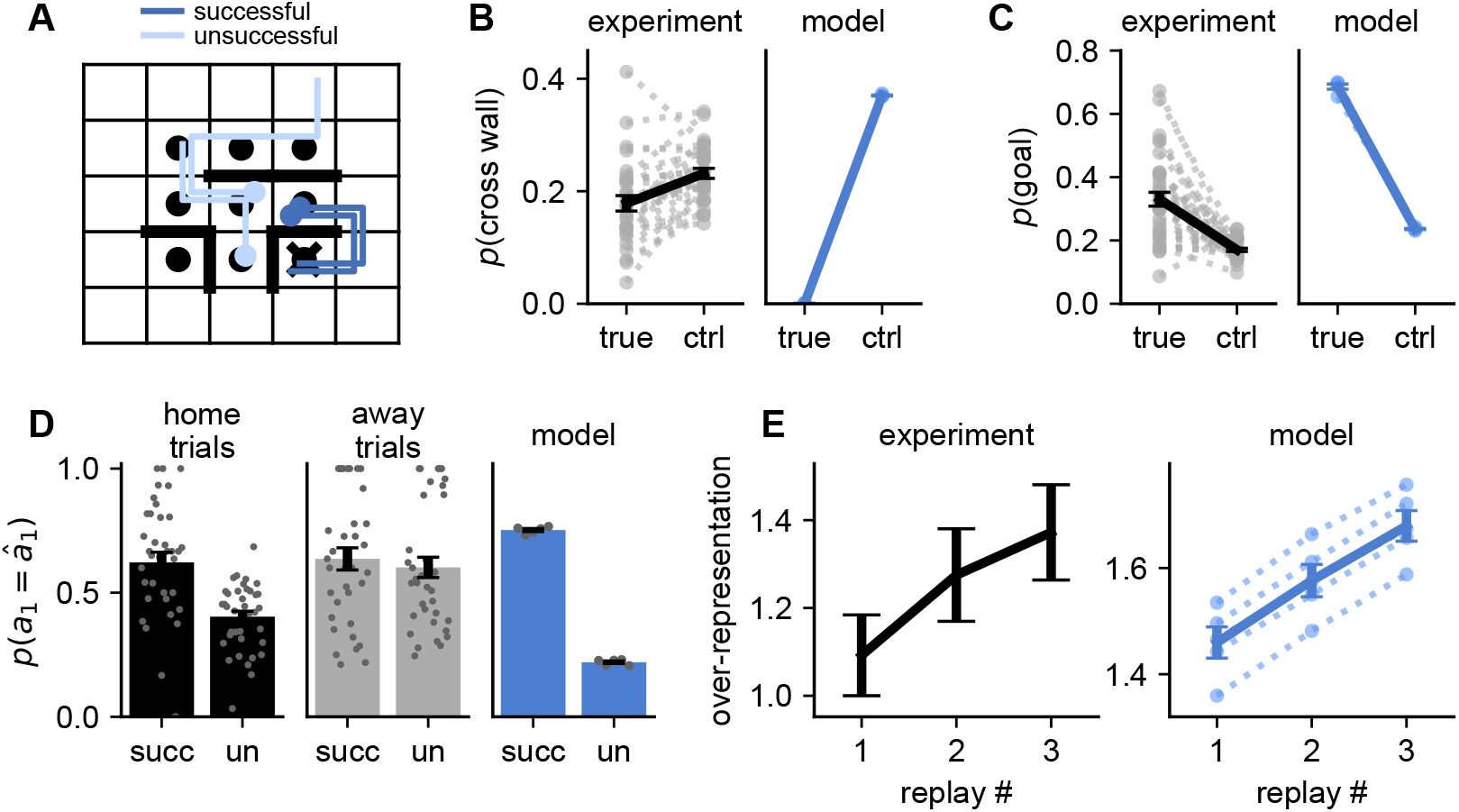
Hippocampal replays resemble model rollouts. **(A)** Illustration of experimental task structure and example replays (Widloski and Foster, 2022). Each episode had a different wall configuration and a randomly sampled home location (cross). Between each ‘home’ trial, the animal had to move to an ‘away’ location, which was sampled anew on each trial (black circles). Colored lines indicate example replay trajectories originating at the blue dots. Replays were detected during the stationary periods at the away locations before returning to the home location and classified according to whether they reached the home location (dark vs. light blue lines). **(B)** Fraction of replay transitions that pass through a wall in the experimental (black) and model (blue) data. Control values indicate the fraction of wall crossings in resampled environments with different wall configurations. Dashed lines indicate individual biological sessions or RL agents, and solid lines indicate mean and standard error across sessions or RL agents. **(C)** Fraction of replays that pass through the goal location in experimental (black) and model (blue) data. Control values indicate the average fraction of replays passing through a randomly sampled non-goal location (Methods). Dashed and solid lines are as in (B). There was no effect for the away trials, where the goal was unknown (Figure S12). **(D)** Probability of taking the first replayed action, *p*(*a*_1_ = *â*_1_), for successful and unsuccessful replays during home trials (left; black), away trials (center; gray), and in the RL agent (right; blue). Bars and error bars indicate mean and standard error across sessions or RL agents (gray dots). **(E)** Over-representation of successful replays during trials with at least three replays in the experimental data (left) and RL agents (right). The over-representation increased with replay number; an effect not seen in the away trials (Figure S12). Over-representation was computed by dividing the success frequency by a reference frequency computed for randomly sampled alternative goal locations. Bars and error bars indicate mean and standard error across replays pooled from all animals (left) or standard error across five RL agents (right; dashed lines).

Similar to previous work (Widloski and Foster, 2022), we found that hippocampal replays avoided passing through walls to a greater extent than expected by chance (Figure 4B; *p <* 0.001, permutation test). This finding suggests that hippocampal replays are shaped by a rapidly updating internal model of the environment, similar to how forward rollouts in the RL agent are shaped by its internal world model (Figure 1A). Additionally, the goal location was over-represented in the hippocampal replays, consistent with the assumption of on-policy rollouts in the RL agent (Figure 4C; *p <* 0.001, permutation test; Widloski and Foster, 2022).

Inspired by our findings in the RL agent, we investigated whether a replayed action was more likely to be taken by the animal if the replay was successful than if it was unsuccessful. Here, we defined a ‘successful’ replay as one which reached the goal location without passing through a wall (Figure 4A). Consistent with the RL model, the first simulated action in biological replays agreed with the next physical action more often for successful replays than for unsuccessful replays (Figure 4D, black; *p <* 0.001, permutation test). Such an effect was not observed in the ‘away’ trials (Figure 4D, gray; *p* = 0.129, permutation test), where the animals had no knowledge of the reward location and therefore could not know what constituted a successful replay. These findings are consistent with the hypothesis that successful replays should increase the probability of taking the replayed action, while unsuccessful replays should decrease this probability.

In the RL agent, we have direct access to the momentary policy and could therefore quantify the causal effect of a replay on behavior (Figure 3E). In the biological circuit, it is unknown whether the increased probability of following the first action of a successful replay is because the replay altered the policy (as in the RL agent), or whether the replay reflects a baseline policy that was already more likely to reach the goal prior to the replay. To circumvent this confound, we analyzed consecutive replays while the animal remained stationary. If our hypotheses hold, that (i) hippocampal replays resemble on-policy rollouts of an imagined action sequence, and (ii) performing a replay improves the policy, then consecutive replays should become increasingly successful even *in the absence* of any behavior between the replays.

To test this prediction, we considered trials where the animal performed a sequence of at least 3 replays at the ‘away’ location before moving to the ‘home’ location. We then quantified the fraction of replays that were successful as a function of the replay index within the sequence, after regressing out the effect of time (Methods; Ólafsdóttir et al., 2017). We expressed this quantity as the degree to which the true goal was over-represented in the replay events by dividing the fraction of successful replays by a baseline calculated from the remaining non-goal locations, such that an over-representation of 1 implies that a replay was no more likely to be successful than expected by chance. This over-representation increased with each consecutive replay during the home trials (Figure 4E; left), and both the second and third replays exhibited substantially higher over-representation than the first replay (*p* = 0.068 and *p* = 0.009 respectively; permutation test; Methods). Such an effect was not seen during the away trials, where the rewarded location was not known to the animal (Figure S12).

These findings are consistent with a theory in which replays represent on-policy rollouts that are used to iteratively refine the agent’s policy, which in turn improves the quality of future replays – a phenomenon also observed in the RL agent (Figure 4E, right). In the RL agent, this effect could arise in part because the agent is less likely to perform an additional rollout after a successful rollout than after an unsuccessful rollout (Figure S13). To eliminate this confound, we drew two samples from the policy each time the agent chose to perform a rollout, and we used one sample to update the hidden state of the agent, while the second sample was used to compute the goal over-representation (Methods). Such decoupling is not feasible in the experimental data, since we cannot read out the ‘policy’ of the animal. This leaves open the possibility that the increased goal over-representation with consecutive biological replays is in part due to a reduced probability of performing an additional replay after a successful replay. However, we note that (i) the rodent task was not a ‘reaction time task’, since a 5-15 s delay was imposed between the end of reward consumption and the next reward becoming available. This makes a causal effect of replay success on the total number of replays less likely. (ii) if such an effect does exist, that is also consistent with a theory where hippocampal replays guide planning.

### How do rollouts improve behavior?

We have now seen that both biological and artificial agents appear to use policy rollouts to improve behavior based on the content of the rollouts. Since all weights of the RL agent are fixed at test time, the only way to achieve this is through optimization of its hidden state. Such an optimization process requires the agent to approximate the gradient of the expected reward with respect to its hidden state, using information from the rollout. To understand how this is possible, we recall that optimization of the weights of the network at the outer level of training uses precisely such policy rollouts as part of a policy gradient algorithm to approximate the gradient of the expected reward with respect to the network parameters. These algorithms consider putative on-policy action sequences *τ* and apply *parameter updates* that cause *p*(*τ*) to increase under the agent’s policy if *τ* led to more reward than expected, and to decrease otherwise (Jensen, 2023b). In our trained agent, adaptation to each new maze does not involve modifications of the fixed network parameters but instead occurs through changes to the *hidden state* ***h***_*k*_. The performance improvements resulting from policy rollouts (Figure 3A; Figure 4E) can therefore be achieved through iterative modifications of ***h***_*k*_ that approximate policy gradient ascent on the expected future reward in the episode as a function of ***h***_*k*_ (Figure 5A). Importantly, while the optimization at the outer loop was hardcoded, this ‘inner’ policy gradient algorithm, operating on the basis of imagined experience, has to be meta-learned by the agent itself over the course of many episodes of the navigation task.

**Figure 5:**
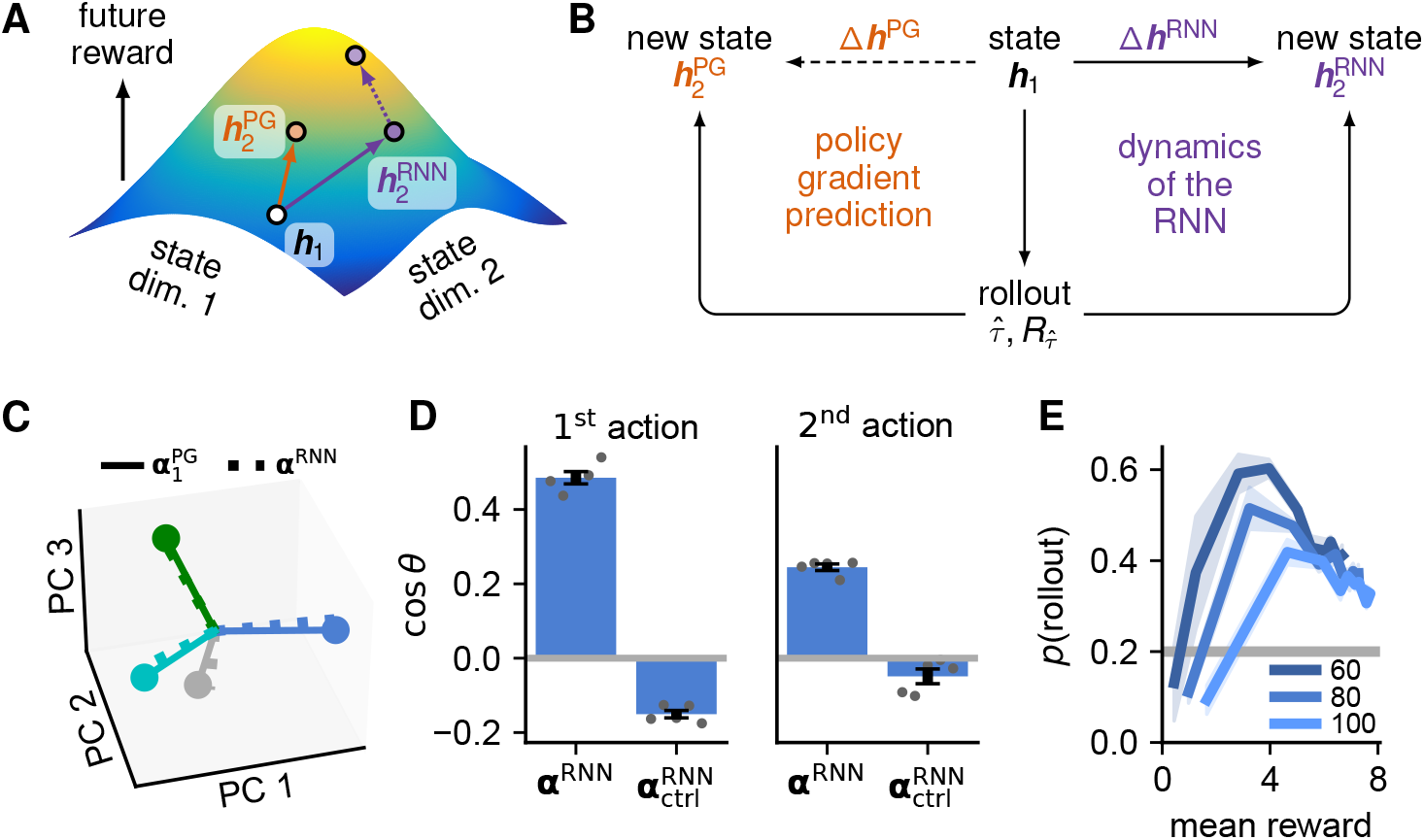
Rollouts implement a hidden state optimization. **(A)** The hidden state ***h***_*k*_ of the RNN induces a policy with an expected future reward for the current episode, setting the ‘initial state’ of a dynamical system consisting of both the agent itself and the environment. Rollouts can improve performance by shifting ***h*** to a region of state space with higher reward for a given environment and agent location 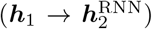. The policy gradient algorithm uses samples from the policy to estimate the direction of steepest ascent of the expected reward as a function of 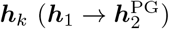. **(B)** We compare this theoretical hidden state update 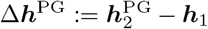 to the empirical hidden state update 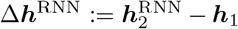 actually performed by the network dynamics on the basis of a rollout 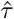 and its associated reward 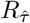. **(C)** A latent space was defined by performing PCA on 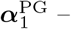 the effect of 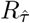 on ***h***_*k*_ under the policy gradient algorithm. Solid lines and circles indicate the normalized average 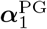 for each of the four possible simulated actions (*â*_1_; colors). Dashed lines indicate the normalized average values of ***α***^RNN^ for the corresponding actions, which are aligned with 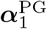 in accordance with the theory. The first 3 PCs capture 100% of the variance in 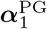, since the policy is normalized and therefore only has three degrees of freedom. **(D)** Average cosine similarity between ***α***^RNN^ and 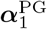, quantified in the space spanned by the top 3 PCs of 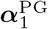. ***α***^RNN^ was computed using the true input, while 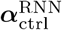 was computed after altering the feedback from the rollout to falsely inform the agent that it had simulated a different action *â*_1,ctrl_ ≠ *â*_1_. Left panel considers the effect of 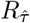 on the first action 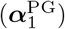 and right panel considers the effect of 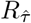 on the second action 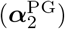. Error bars indicate standard error across five RL agents (gray dots). **(E)** We trained networks of different sizes (legend) and quantified both their performance (x-axis) and frequency of performing a rollout (y-axis) over the course of training (Figure S3). To reach a given performance, smaller networks relied more on rollouts, suggesting that the RL agents learn to plan in part because they are capacity limited. Agents initially did not perform frequent rollouts, as they had not learned a world model or how to integrate rollouts with the recurrent policy. This was followed by an increase in rollout frequency to substantially above chance levels, followed by a gradual transition to fewer rollouts late in training as the agents became increasingly good at the task, suggesting that they also plan because they are data limited. This is consistent with human behavior, where response times decreased over the course of meta-learning (Figure S3B).

To verify that the algorithm implemented by the agent approximates such single-sample policy gradient estimates, we considered each rollout performed by the RL agent and computed both (i) the actual hidden state update performed on the basis of this rollout, and (ii) the expected hidden state update computed by applying the policy gradient algorithm to the same rollout (Figure 5B; Methods). The policy gradient algorithm specifies that rollouts should change ***h***_*k*_ in a way that increases 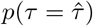 if the rollout is better than some baseline and decreases 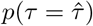 otherwise. Since we do not know the baseline, we performed our analysis by taking the *derivative* of the hidden state change with respect to the expected reward from physically following 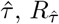, which is independent of the baseline (Methods). This allowed us to define (i) a quantity 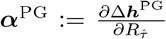 that predicts how the hidden state *should* change as a function of 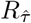 in the policy gradient formulation, and (ii) the corresponding quantity 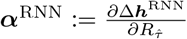 that indicates how the hidden state *actually* changed as a function of the content of the rollout. If the agent performs approximate policy gradient ascent in hidden state space, ***α***^RNN^ should be aligned with ***α***^PG^.

We began by considering the effect of the first action in the rollout, *â*_1_, on the hidden state of the agent. We did this by querying the alignment between (i) ***α***^RNN^ computed across rollouts from 1,000 episodes, and (ii) 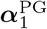 computed from the same rollouts when considering only the probability of executing *â*_1_. To visualize this alignment, we performed PCA on 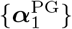 from all rollouts and projected both 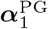 and ***α***^RNN^ into this low-dimensional subspace. We then computed the average of each of these two quantities for each simulated action *â*_1_ *∈ {*left, right, up, down*}* . The average value of ***α***^RNN^ was strongly aligned with the average value of 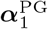 for each action (Figure 5C), consistent with the policy gradient algorithm. This implies that 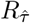 has *different* effects on the policy depending on the replayed trajectory 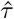. In other words, the spatial content of the rollout dynamically modulates the way in which the reward signal from the rollout affects the hidden state and policy of the agent.

To quantify the overlap between ***α***^RNN^ and 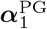 on a rollout-by-rollout basis, we computed the average cosine similarity *d* between ***α***^RNN^ and 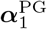 across all rollouts. This overlap was substantially larger than zero (*d* = 0.49 *±*0.02 mean*±* sem; Figure 5D, left). When instead computing the overlap with 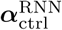 after changing the feedback input to falsely inform the agent that it simulated a different action *â*_1,ctrl_ *≠â*_1_, the similarity was *d* = *−*0.15 *±* 0.01. This confirms that ***h***_*k*_ is optimized by incorporating the specific feedback input obtained from the rollout, and the negative sign reflects anti-correlations due to the policy being a normalized distribution over actions. For these analyses, we only considered the first simulated action *â*_1_. When instead querying the effect of the rollout on subsequent actions in 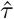, we found that the feedback input was also propagated through the network dynamics to these later actions, although with a weaker alignment than for the first action (Figure 5D, right). This weaker effect for the second action could arise because the agent has the capacity to re-plan at each iteration, which is not taken into account in the rollouts, or it could simply reflect additional noise resulting from the network dynamics.

## Discussion

We have developed a new theory of planning in the prefrontal-hippocampal network, implemented as a recurrent neural network model and instantiated in a spatial navigation task requiring multi-step planning (Figure 1). This model consists of a recurrent meta-reinforcement learning agent augmented with the ability to plan using policy rollouts, and it explains the structure observed in human behavior (Figure 2). Our results suggest that mental rollouts could play a major role in the striking human ability to adapt rapidly to new tasks by facilitating behavioral optimization without the potential cost of executing suboptimal actions. Since mental simulation is generally faster and less risky than physical actions (Vul et al., 2014), this can improve overall performance despite the temporal opportunity cost of simulation (Figure 3; Agrawal et al., 2022; Hamrick et al., 2017).

Our theory also suggests a role for hippocampal replays during sequential decision making. A reanalysis of rat hippocampal recordings during a navigation task showed that patterns of hippocampal replays and their relationship to behavior resembled those of rollouts in our model (Figure 4). These results suggest that hippocampal forward replays could be a core component of planning, and that the mechanistic insights derived from our model could generalize to biological circuits. In particular, we hypothesize that forward replays should affect behavior differently depending on whether they lead to high-value or low-value states (Figure 3; Wu et al., 2017). This hypothesis is consistent with previous models where hippocampal replay is used to update state-action values that shape future behavior (Mattar and Daw, 2018). We suggest that forward replay implements planning through feed-back to prefrontal cortex that drives a ‘hidden state optimization’ reminiscent of recent models of motor preparation (Figure 5; Kao et al., 2021). Such model-based policy refinement differs from some prior work in the reinforcement learning literature that involves arbitration between model-free and model-based policies computed separately (Daw et al., 2005; Geerts et al., 2020). Instead, model-based computations in our framework iteratively update a single policy that can be used for decision making at different stages of refinement. This is consistent with previous work proposing that model-based computations are used to iteratively refine values or world models learned via a model-free mechanisms (Keramati et al., 2016; Momennejad et al., 2017).

### Neural mechanisms of planning and decision making

Our model raises several interesting hypotheses about neural dynamics in hippocampus and prefrontal cortex and how these dynamics affect behavior. One is that hippocampal replays should causally affect the behavior of an animal as also suggested in previous work (Foster, 2017; Pfeiffer and Foster, 2013; Widloski and Foster, 2022). However, this has been difficult to test experimentally due to the confound of how the behavioral intentions of the animal itself affect the content of hippocampal replays (Foster, 2017). Perhaps more interestingly, we predict that hippocampal replays should directly affect PFC representations, consistent with previous work showing coordinated activity between hippocampus and PFC during sharp-wave ripples (Jadhav et al., 2016). Specifically, PFC activity should change to make replayed actions more likely if the replayed trajectory is better than expected and less likely if worse than expected. These predictions can be investigated in experiments that record neural activity simultaneously from hippocampus and PFC, where both the timing and qualitative change in PFC representations can be related to hippocampal replays. This would be most natural in rodent experiments with electrophysiology, although human experiments using MEG for replay detection could potentially also investigate the effect of replays on cortical representations and behavior (Kurth-Nelson et al., 2016; Liu et al., 2019, 2021).

While we propose a role of hippocampal replays in shaping immediate behavior via recurrent network dynamics, this is compatible with replays also having other functions over longer timescales, such as memory consolidation (van de Ven et al., 2016; Carr et al., 2011) or dopamine-driven synaptic plasticity (Gomperts et al., 2015; De Lavilléon et al., 2015). Additionally, we only considered the case of local forward replays and showed that they can be used to drive improved decision making. These replays will always have high ‘need’ according to the theory of Mattar and Daw (2018), since they start at the current agent location and visit likely upcoming states. They should similarly have a high ‘gain’, since rollouts lead to an increase in expected future reward (Figure 3A). However, our choice to focus on local replays in this work does not imply that non-local or reverse replays could not play a similar role. Backwards planning from a goal location is for example more efficient in environments where the branching factor is larger in the forward than the reverse direction, and branching ‘rollouts’ have been shown to improve performance in previous RL models (Pascanu et al., 2017).

### Hippocampal replay and theta sequences

This work focuses on the trade-off between ‘thinking’ and acting, investigating internal computations that can improve decision making without additional physical experience. This is the phenomenon investigated in human behavior, where the analyses focused on stopping times. It is also an explicit feature of the RL agent, which chooses between acting and performing a rollout rather than doing both simultaneously. A putative neural correlate of such planning in the absence of behavior is hippocampal replay, given the ubiquitous finding that it occurs primarily when animals are stationary (Foster, 2017) and its hypothesized role in decision making (Pfeiffer and Foster, 2013; Widloski and Foster, 2022; Foster, 2017; Mattar and Daw, 2018). While some have challenged these ideas (Papale et al., 2016; Carey et al., 2019; Gillespie et al., 2021), our analyses show that a replay-like mechanism could in principle drive improved decision making in a manner consistent with human behavior.

Another phenomenon suggested to play a role in decision making is that of hippocampal theta sequences (Johnson and Redish, 2007; Wikenheiser and Redish, 2015a,b). Theta sequences typically represent states in front of the animal (Johnson and Redish, 2007) and are affected by the current goal location (Wikenheiser and Redish, 2015b), similar to what we found in our analyses of hippocampal replays (Figure 4). However, since theta sequences predominantly occur during active behavior, they are potentially less relevant than hippocampal replays for the tradeoff between acting and thinking. Nonetheless, our RL model also suggests a potential mechanism by which short theta sequences could guide behavior by providing recurrent feedback to cortical decision making systems about immediately upcoming states and decision points. This mechanism would likely require feedback in the form of estimated value functions rather than predicted empirical reward, since shorter rollouts are less likely to visit ‘terminal’ rewarding states than longer-reaching replays. Under this hypothesis, hippocampal replays could provide a mechanism for longer-term planning during stationary periods, while theta sequences would help modify these plans on-the-go via short-term predictions – both operating through recurrent feedback to frontal cortex.

### Alternative planning algorithms

Planning in the RL agent was carried out explicitly in the space of observations. While this was already an abstract representation rather than pixel-level input, it could be interesting to explore planning in a latent space optimized e.g. to predict future observations (Zintgraf et al., 2019) or future policies and value functions (Ho et al., 2022; Schrittwieser et al., 2020). These ideas have proven useful in the machine learning literature, where they allow models to ignore details of the environment not needed to make good decisions, and it is plausible that the internal model of humans similarly does not include such task-irrelevant details. We also assumed that the planning process itself was ‘on policy’ – that is, the policy that was used to sample actions in the planning loop was identical to the policy used to act in the world. Although there is some support from the hippocampal replay data that forward replays are related to the ‘policy’ (e.g. wall avoidance and goal over-representation; Figure 4), there is in theory nothing that prevents the planning policy from differing arbitrarily from the action policy. In fact, the planning policy could even be explicitly optimized to yield good *plans* rather than re-using a policy optimized to yield good *behavior* (Pascanu et al., 2017). Such off-policy hippocampal sequence generation has also formed the basis of other recent theories of the role of hippocampus in planning and decision making (McNamee et al., 2021; Mattar and Daw, 2018). In this case, the policy gradient view of rollouts still provides a natural language for formalizing the planning process, since numerous off-policy extensions of the canonical policy gradient algorithm exist (Peshkin and Shelton, 2002; Jie and Abbeel, 2010).

### Why do we spend time thinking?

Despite our results showing that humans and RL agents make extensive use of planning, mental simulation does not generate fundamentally new information. In theory, it should therefore be possible to make equally good ‘reflexive’ decisions given enough experience and computational power. However, previous work has shown that imagination can affect human choices (Gershman et al., 2017), and that having more time to process the available information can improve decisions (Gershman et al., 2014). This raises questions about the computational mechanism whereby choices are modulated by such a time-consuming process and why decision making often takes time rather than being instantaneous. One possible reason could be that our decision making system is capacity limited and does not have enough computational power to generate the optimal policy (Russek et al., 2022). In our computational model, this is supported by the observation that agents consisting of smaller RNNs tend to perform more rollouts than larger agents (Figure 5E). Alternatively, we could be data limited, meaning that we have not received enough training to learn the optimal policy. This also has support in our computational model, where networks of all sizes perform many rollouts early in training, when they have only seen a small amount of data, and gradually transition to a more reflexive policy that relies less on rollouts (Figure 5E; Figure S3).

We hypothesize that data limitations are a major reason for the use of temporally extended planning in animals. In particular, we reason that learning the instantaneous mapping from states to actions needed for reflexive decisions would require a prohibitive amount of training data, which is generally not available for real-life scenarios. Indeed, training our meta-reinforcement learner required millions of episodes, while humans performed well immediately after seeing a simple description and demonstration (Figure S3). Such rapid learning could be due in part to the use of temporally extended planning algorithms as a form of ‘canonical computation’ that generalizes across tasks. If this is the case, we would be able to rely on generic planning algorithms acquired over the course of many previous tasks in order to solve a new task. When combined with a new task-specific transition function learned from relatively little experience or inferred from sensory inputs, planning would facilitate data-efficient reinforcement learning by trading off processing time for a better policy (Schrittwieser et al., 2020). This is in contrast to our current model, which had to learn from scratch both the structure of the environment *and* how to use rollouts to shape its behavior. Importantly, planning as a canonical computation could generalize not just to other navigation tasks but also to other domains, such as compositional reasoning and sequence learning, where replay has recently been demonstrated in humans (Liu et al., 2019; Schwartenbeck et al., 2023; Liu et al., 2021).

## Author contributions

KTJ, GH, and MM conceived of the project and developed the human experimental paradigm. KTJ performed all simulations and analyzed the data. All authors interpreted the results and wrote the paper.

## Acknowledgments

We are grateful to Widloski and Foster (2022) for sharing their data with us, to Ta-Chu Kao for assistance with development of the software library used to train and analyze RL agents, and to Wei Ji Ma for suggesting the method used to estimate human thinking times. We thank Nathaniel Daw, Wei Ji Ma, Zeb Kurth-Nelson, Thomas Akam, Karen Schroeder, John Widloski, Jake Stroud, Fred Callaway, Ta-Chu Kao, Evan Russek, Marine Schimel, Ji-An Li, Jeroen Olieslagers, and Sixuan Chen for helpful feedback on the manuscript. KTJ was funded by a Gates Cambridge scholarship. For the purpose of open access, the authors have applied a Creative Commons Attribution (CC BY) licence to any Author Accepted Manuscript version arising from this submission.

## Data availability

Human behavioral data is available on github. For the rodent data, we refer to Widloski and Foster (2022).

## Code availability

Code for model training and all analyses is available on github.

## Supplementary figures

**Figure S1:**
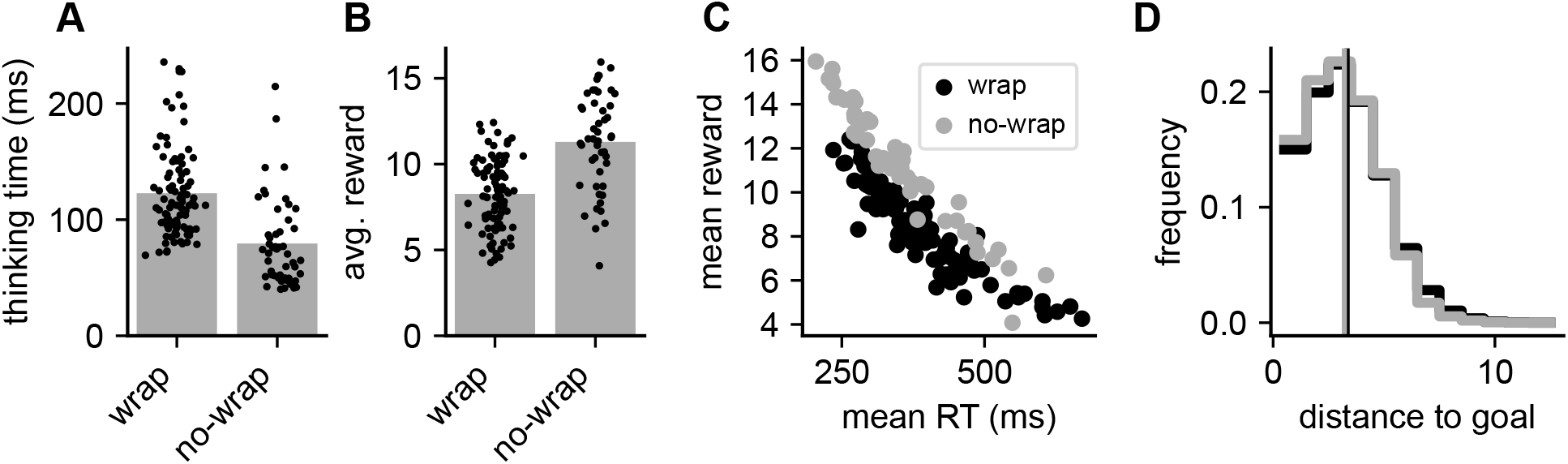
Comparison of human behavior in mazes with periodic and non-periodic boundaries. For comparison with the data collected with periodic maze edges, we collected data from 49 additional participants performing the same task in non-periodic mazes. These non-periodic mazes were generated such that the average shortest path length was similar to the periodic mazes considered in all other analyses (Methods). **(A)** Mean across participants (black dots) of the average thinking time during non-guided exploitation trials. This is computed separately for participants performing the task with periodic (left) or non-periodic (right) boundaries. The average thinking time was significantly higher with periodic boundaries (*p <* 0.001; permutation test). **(B)** As in (A), now quantifying average reward per episode across participants. The average reward was significantly higher for participants with non-periodic boundaries than participants with periodic boundaries (*p <* 0.001; permutation test). **(C)** Scatter plots of average reward against average response time (across all actions) for participants with periodic (black) or non-periodic (gray) boundaries. For a given average response time, participants with non-periodic boundaries get more reward. The results in (A)-(C) are all consistent with participants having a worse ‘model-free’ policy in the unfamiliar environment with periodic boundaries, which leads to more thinking and worse performance for a given amount of thinking. **(D)** Histogram of distances to reward for randomly selected start locations in randomly generated mazes with either periodic (black) or non-periodic (gray) boundaries. Vertical lines indicate the mean distance to reward in each setting. The distributions of minimum path lengths are closely matched across settings, suggesting that different path lengths are not the reason for the effects observed in (A)-(C).

**Figure S2:**
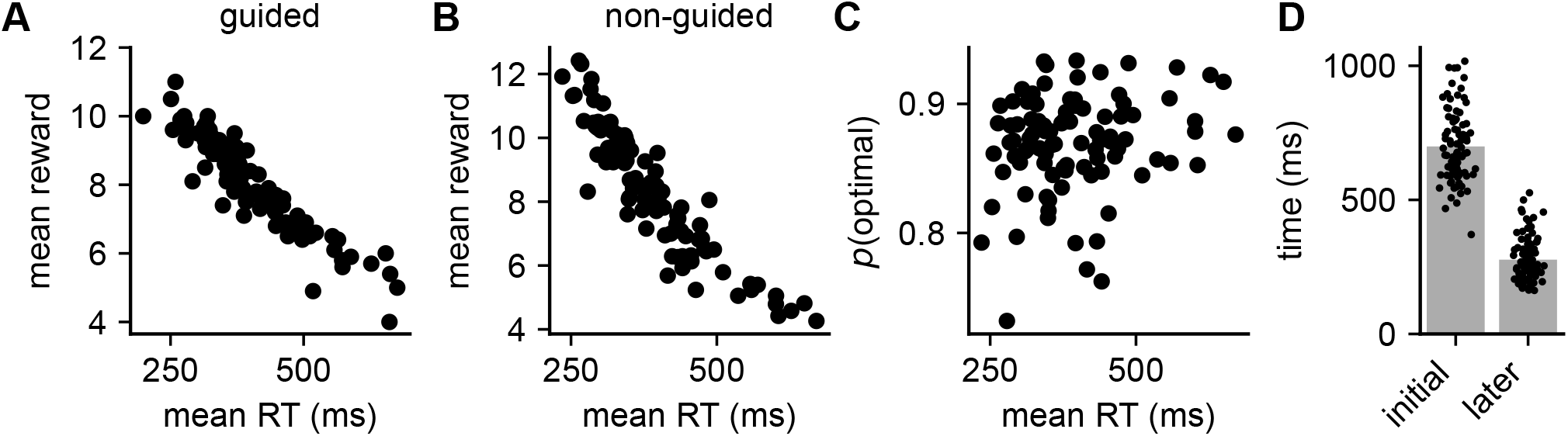
Overview of human data for all participants. **(A)** Mean reward per episode as a function of the average response time during the guided trials (Methods). Each data point corresponds to a single participant. **(B)** Mean reward per episode as a function of the average response time during the non-guided trials. The strong negative correlation implies that participants on average got more reward when they acted faster, confirming that participants who acted faster were not simply making random key presses. **(C)** Fraction of actions that were consistent with an optimal policy as a function of mean response time, plotted across participants (black dots) during the non-guided trials. There was a significant positive Pearson correlation between these two quantities (*r* = 0.20; *p* = 0.024, permutation test). This correlation confirms that participants who thought for longer were not simply disengaged with the task, but that they instead invested the time to make higher-quality decisions. One outlier with *p*(optimal) = 0.45 was excluded from this analysis. **(D)** Mean of the log-normal distribution of perception-action delays fitted to data from the guided episodes for each participant (dots) using either the first action within each trial (left) or all other actions (right). These prior distributions were used to infer the thinking times in Figure 2.

**Figure S3:**
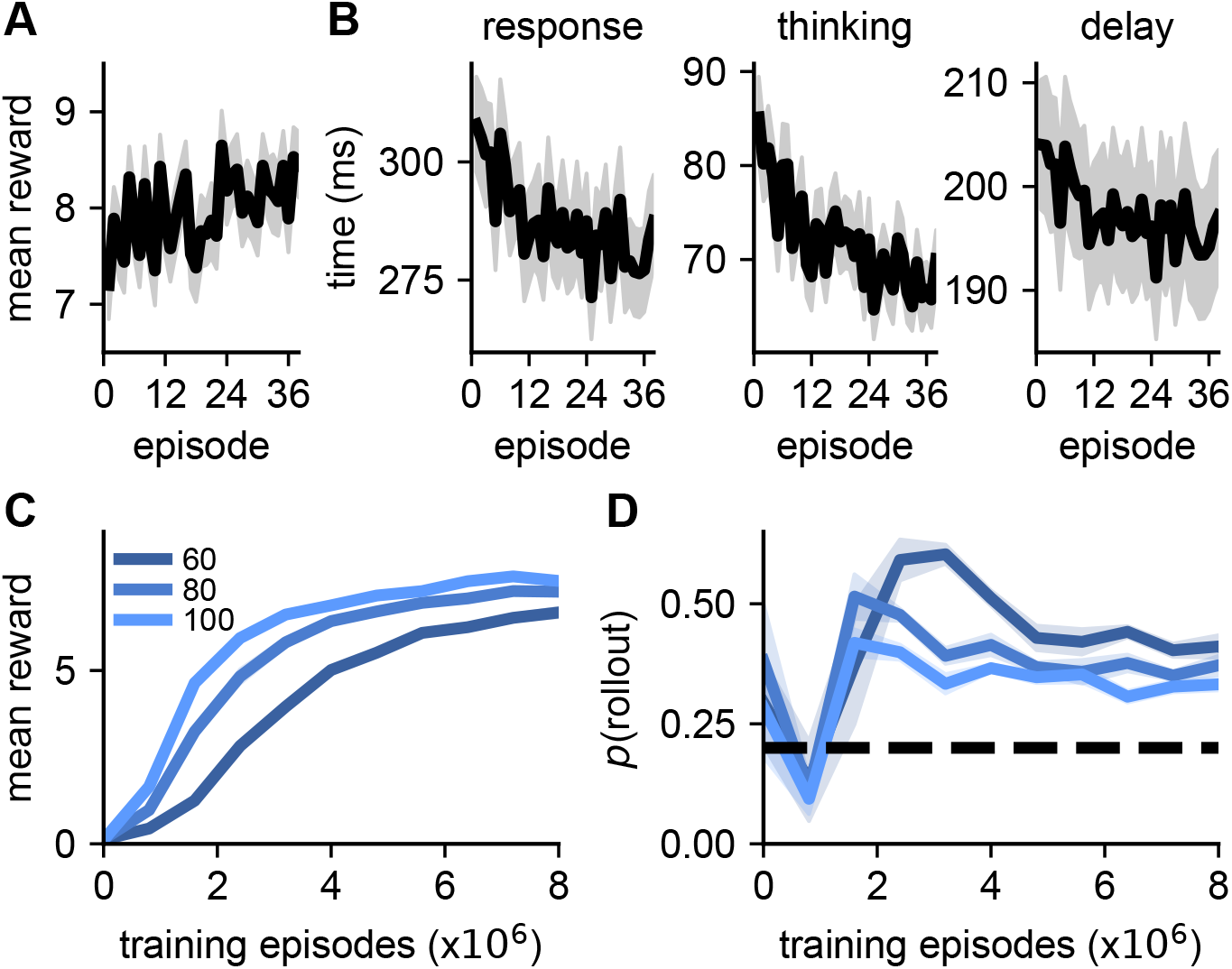
Performance and response times over learning in humans and RL agents. **(A)** Mean and standard error across human participants of the average reward per episode over the course of the 38 episodes used for all analyses. There was a positive correlation between episode number and average reward of *r* = 0.075*±* 0.021 (mean ± sem) across participants. **(B)** Mean and standard error across human participants of the median time per action for each participant over the course of the 38 episodes used for all analyses. We plot this as both the total response time (left) as well as the thinking time (center) and sensorimotor delay (right) for each action (see Methods for details of how these are computed). We observed a negative correlation of *r* = *−*0.107 *±*0.019 (mean*±* sem) across participants between episode number and median response time, *r* = *−*0.146 *±*0.021 for median thinking time, and *r* = *−*0.114 *±*0.019 for median sensorimotor delay. The decrease in average median response time from the first five to the last five episodes is 6.8%, while the increase in average reward per episode is 7.6%, suggesting that a substantial part of the increase in reward could be due to faster decision making. **(C)** We trained networks of different sizes (legend; *N∈ {*60, 80, 100 *}*) and quantified their performance over the course of training. **(D)** Fraction of timesteps where the agent chose to perform a rollout over the course of training for different network sizes. The agents perform rollouts at chance level but with high variance at initialization, and this data point was therefore not included in the analysis in Figure 5E, where we only considered the learned rollout frequency from episode 800,000 onwards. The agents first learn to suppress the rollout frequency below chance before increasing it to levels above chance. This is consistent with a theory where rollouts only become useful when (i) an internal world model has been learned, and (ii) the agent has learned how to use rollouts to improve its policy. Finally, rollouts become less frequent again later in training as the base policy improves, which is similar to how humans become faster across episodes. We hypothesize that humans start in this regime from episode 1 because they (i) construct a mental ‘world model’ immediately upon seeing the task, and (ii) already know how to integrate planning with decision making.

**Figure S4:**
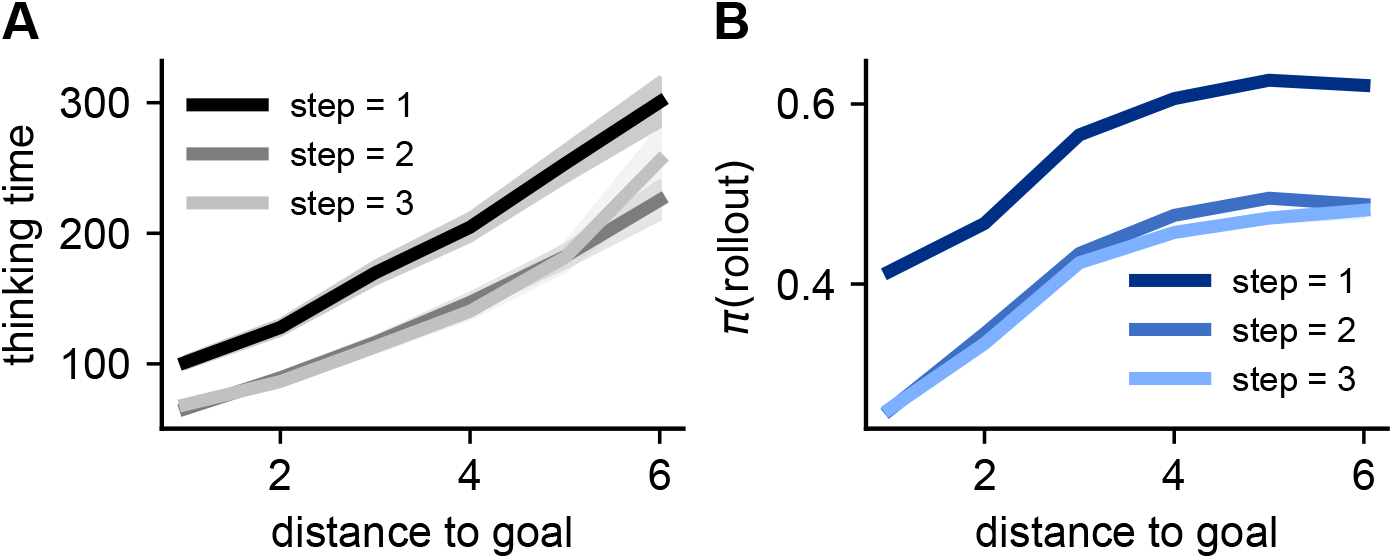
Thinking time and *π*(rollout) by momentary distance to goal and step within trial. **(A)** Figure illustrating the average thinking time across human participants as a function of momentary distance-to-goal (x-axis), conditioned on different steps within the trial (lines, legend). Subjects generally spent longer thinking before the first action of each trial, after controlling for the momentary distance-to-goal, while subsequent actions were associated with similar thinking times. Lines and shadings indicate mean and standard error when repeating the analysis across human participants (*n* = 94). **(B)** *π*(rollout) for the agent clamped to the human trajectory as a function of momentary distance-to-goal and for different steps within the trial. Similar to the human participants, the agent had a higher probability of performing a rollout on the first step of each trial. Subsequent steps were associated with similar rollout probabilities after controlling for the momentary distance-to-goal. When conditioning on both momentary distance-to-goal and step within the trial, the residual correlation between *π*(rollout) and thinking time remained at a significantly positive value of *r* = 0.026 *±* 0.004 (mean *±* sem).

**Figure S5:**
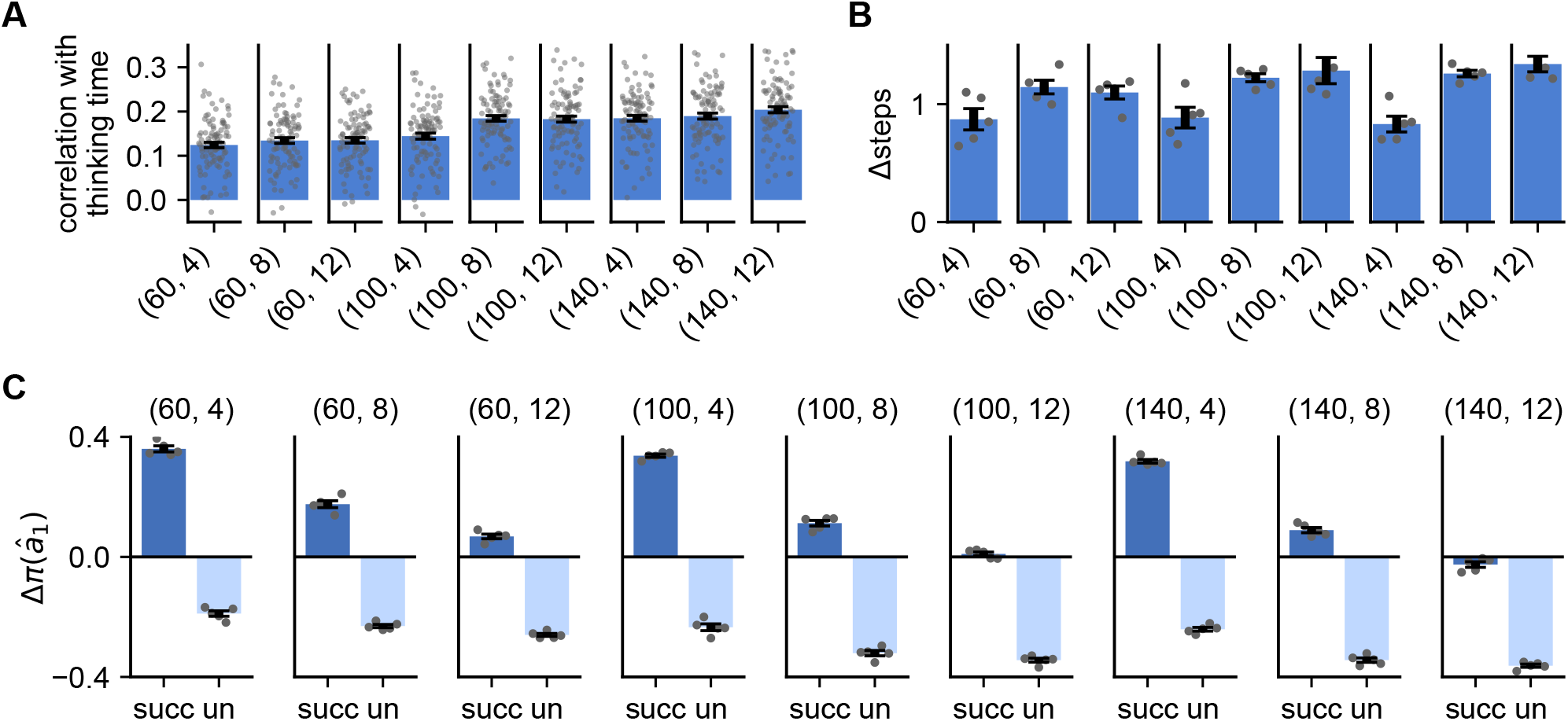
Properties of networks with different hyperparameters. To investigate the robustness of our results to the choice of network size (*N*) and maximum planning horizon (*L*), we trained five networks with each combination of *N ∈ {*60, 100, 140*}* and *L∈ {*4, 8, 12*}* and repeated some of our key analyses. The results in the main text are all reported for a network with *N* = 100 and *L* = 8. **(A)** We quantified the correlation between human response times and the mean *π*(rollout) across five RL agents for each set of hyperparameters (c.f. Figure 2F). x-ticks indicate network size and planning horizon as (*N, L*). Error bars indicate standard error of the mean across human participants (gray dots). **(B)** We computed the improvement in the network policy from performing five rollouts compared to the policy in the absence of rollouts (c.f. Figure 3A). The policy improvement was quantified as the average number of steps needed to reach the goal on trial 2 in the absence of rollouts, minus the average number of steps needed with five rollouts enforced at the beginning of the trial and no rollouts during the rest of the trial. Positive values indicate that rollouts improved the policy. Error bars indicate standard error across five RL agents (gray dots). **(C)** We investigated how rollouts changed the policy (c.f. Figure 3E). For each set of hyperparameters, we computed the average change in *π*(*â*_1_) from before a rollout to after a rollout and report this change separately for successful (‘succ’) and unsuccessful (‘un’) rollouts. Positive values indicate that *â*_1_ became *more* likely and negative values that *â*_1_ became *less* likely after the rollout. Error bars indicate standard error across five RL agents (gray dots). Networks with longer planning horizons tend to have less positive Δ*π*(*â*_1_) for successful rollouts and more negative Δ*π*(*â*_1_) for unsuccessful rollouts. This is consistent with a policy gradient-like algorithm with a baseline that approximates the probability of success, which increases with planning horizon. In other words, since longer rollouts are more likely to reach the goal, we should expect them to be successful and not strongly update our policy when it occurs. On the contrary, an unsuccessful rollout is less likely and should lead to a large policy change. The converse is true for shorter planning horizons.

**Figure S6:**
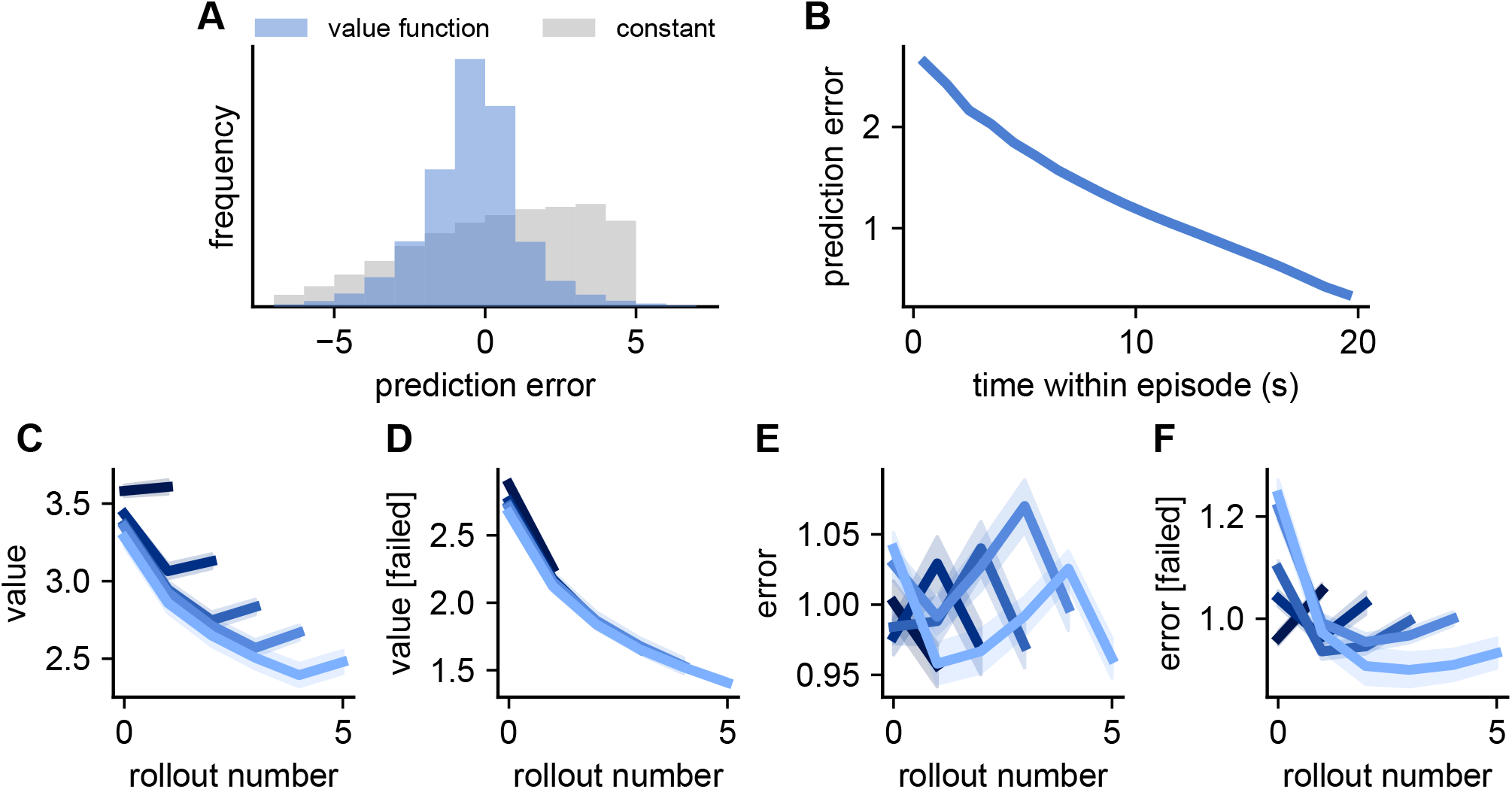
Analyses of RL agent value functions with real and imagined experience. **(A)** Blue histogram indicates the distribution of reward prediction errors across all iterations from five RL agents. The prediction error was defined at each iteration as the value function of the agent minus the true reward-to-go for the rest of the episode. Gray histogram indicates a control computed by subtracting the instantaneous reward-to-go from the mean reward-to-go across all agents and iterations of network dynamics. **(B)** Mean (line) and standard error (shading) across agents of the prediction error as a function of time within an episode. Prediction errors decrease monotonically with time since the agents are required to integrate expected reward over shorter time horizons later in the episode. The rate of decrease is fastest earlier in the episode as the agent initially does not know where the reward is. **(C)** Value function as a function of rollout number. We separated the analysis into sequences of rollouts of different lengths to avoid confounds between the number of rollouts performed and the value of the state. The darkest color corresponds to sequences of a single rollout and the lightest color to sequences of five rollouts, with intermediate colors indicating intermediate numbers of rollouts. Lines and shadings indicate mean and standard error across five RL agents. The value function generally decreases as more rollouts are performed, with an increase in expected reward from the very last rollout. We hypothesized that this final increase in *V* is because successful rollouts are more likely to be followed by a physical action (Figure S13), and that such rollouts are also likely to lead to an increase in expected reward. **(D)** To test this hypothesis, we repeated the analysis in (C), now considering only sequences of rollouts where no rollouts were successful. As expected, the predicted value did not increase for the final rollout in this setting. **(E)** Reward prediction error after a rollout as a function of rollout number. There is a decrease in prediction error after the final rollout, consistent with the increased prevalence of successful rollouts leading to a better estimate of future reward. **(F)** As in (E), now considering only sequences of rollouts where no rollouts were successful. In cases where the agent performed many unsuccessful rollouts, indicating that its policy was bad, the initial value function is also likely to have been.

**Figure S7:**
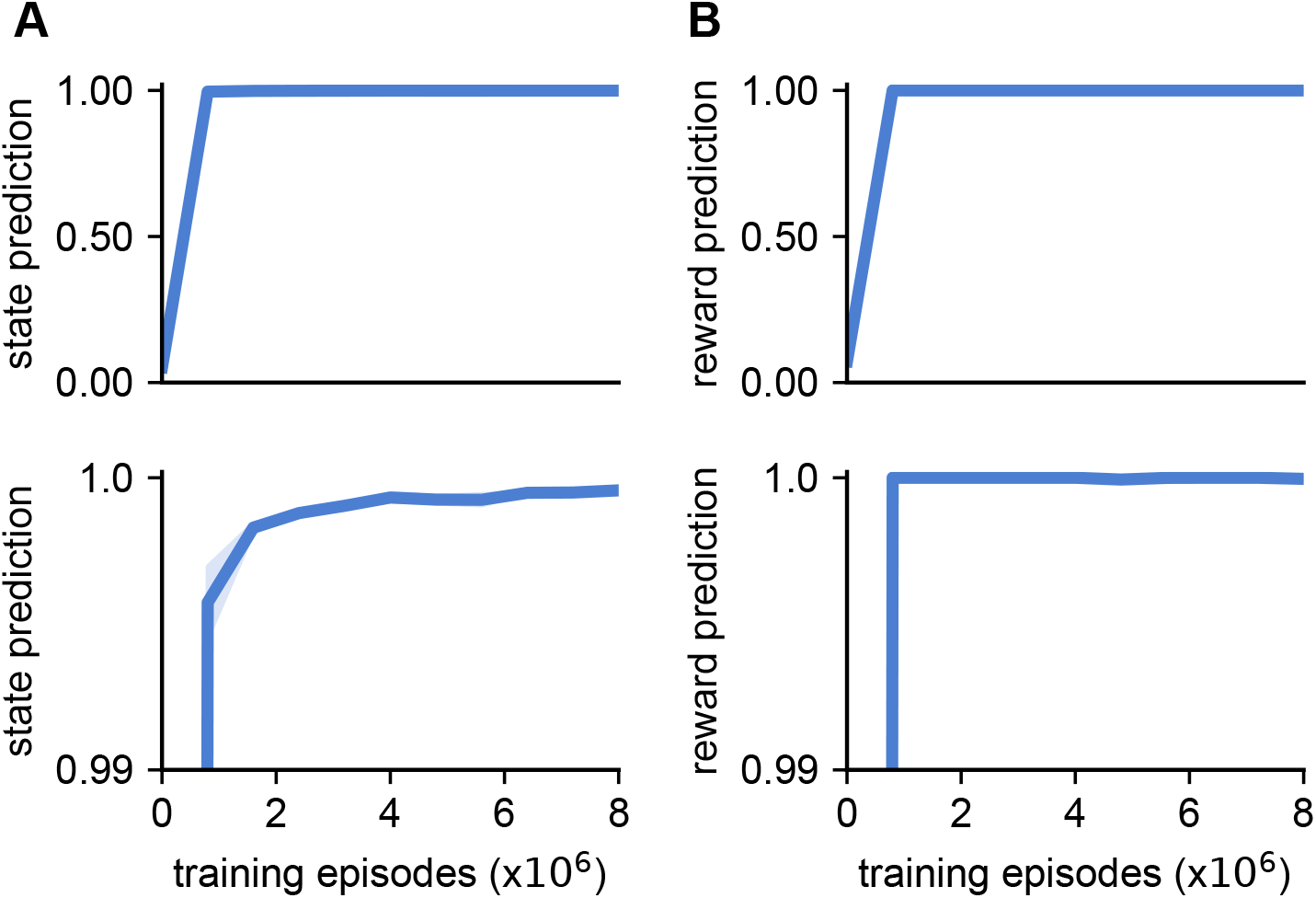
Accuracy of the internal world model. **(A)** Accuracy of the internal transition model over the course of training. Accuracy was computed as the probability that the predicted next state was the true state reached by the agent, ignoring all teleportation steps where the transition cannot be predicted. The accuracy was averaged across all iterations from 1,000 episodes, and the line and shading indicate mean and standard error across five RL agents. The upper panel considers the full range of [0, 1] while the lower panel considers the range [0.99, 1.0]. The transition model rapidly approaches ceiling performance, although it continues to improve slightly throughout training. **(B)** Accuracy of the internal reward model over the course of training. Accuracy was computed as the probability that the predicted reward location was the true reward location during the exploitation phase of the task (see Figure S9 for an analysis of the model accuracy during exploration). Lines and shadings indicate mean and standard error across five RL agents.

**Figure S8:**
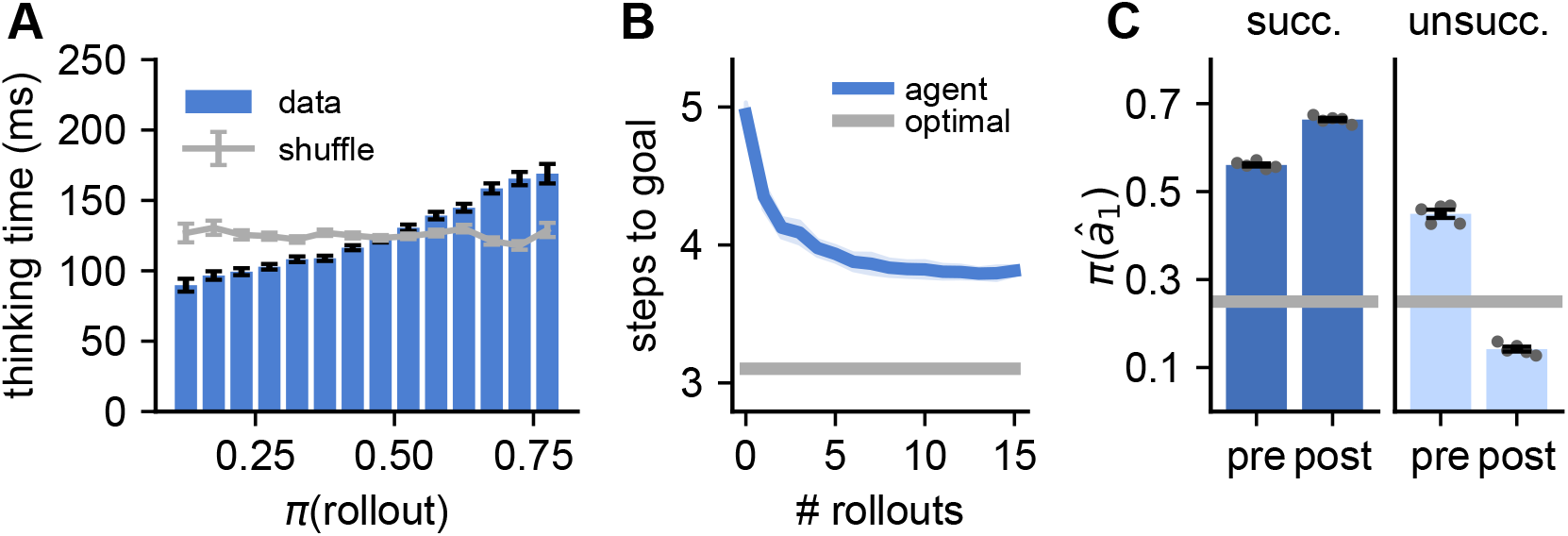
Analyses of RL agents with variable temporal opportunity costs of rollouts. In the main text, RL agents were trained with a constant temporal opportunity cost of 120 ms when performing a rollout, irrespective of the actual rollout length. In this figure, we demonstrate that our main results are not sensitive to this choice by training an additional set of agents with a ‘variable’ rollout time cost of 24 ms per imagined action. This leads to a range of rollout time costs of 24 ms to 192 ms. **(A)** Human thinking time plotted against the probability of the agent performing a rollout (*π*(rollout)) under its policy when exposed to the same mazes and action sequences as the human participants. The correlation between these two quantities was 0.099 *±*0.005 across participants. See Figure 2E for the equivalent plot for agents trained with a constant rollout time cost. **(B)** Average number of physical actions required to reach the goal on trial 2 of an episode as a function of the number of rollouts enforced at the beginning of the episode. See Figure 3A for the equivalent plot for agents trained with a constant rollout time cost. **(C)** Probability of taking the first simulated action of the rollout, *â*_1_, before (*π*^pre^(*â*_1_)) and after (*π*^post^(*â*_1_)) the rollout, evaluated separately for successful (left) and unsuccessful (right) rollouts. See Figure 3E for the equivalent plot for agents trained with a constant rollout time cost.

**Figure S9:**
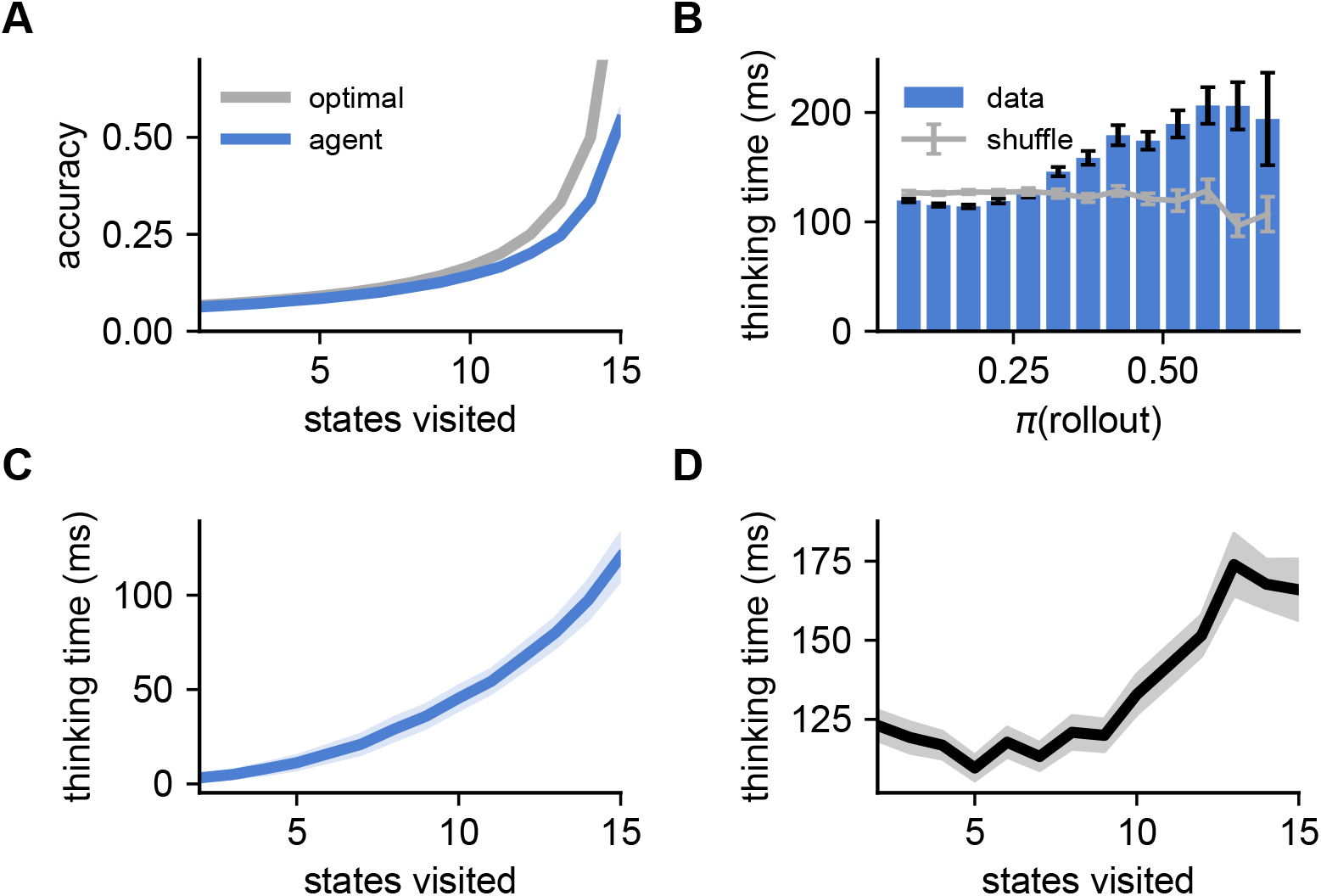
Analyses of the exploration period in humans and RL agents. **(A)** At each iteration, the agent outputs its belief over the goal location under its internal model, which was trained using a cross-entropy loss (Methods). The figure shows the average probability assigned to the true goal, plotted as a function of the number of unique states visited during the exploration phase of the task. As more states are explored, the posterior over possible goals becomes narrower and prediction accuracy increases. When the model chooses to perform a rollout, the imagined goal is chosen as the maximum likelihood location from this posterior to predict the ‘success’ of the rollout. This imagined goal becomes increasingly likely to be the true goal as the agent explores more of the environment. This analysis is consistent with the view of Alver and Precup (2021) that recurrent meta-reinforcement learning agents maintain a ‘belief state’ over the set of tasks they could be in, and that this belief state is gradually updated on the basis of their experience. **(B)** Thinking time of human participants during exploration, plotted as a function of *π*(rollout) for RL agents clamped to the human trajectory. Bars and error bars indicate mean and standard error of the human thinking time across all states where *π*(rollout) fell in the corresponding bin. Gray line indicates a control where human thinking times have been shuffled. The Pearson correlation between *π*(rollout) and human thinking times is *r* = 0.098 *±*0.008, suggesting that the model captures some of the structure in human thinking during exploration and not just during the exploitation phase. The very first action of each episode was not included in this or other analyses of the human data. **(C)** Model thinking time as a function of the number of unique states visited during the exploration phase of the task, with each rollout assumed to take 120 ms as specified in the main text and Methods. Line and shading indicate mean and standard error across RL agents. The increase in thinking time with visited states mirrors the predictive performance from panel (A) and suggests that the agent increasingly chooses to engage in ‘model-based’ planning as its uncertainty over possible goal locations decreases. **(D)** Human thinking time as a function of the number of unique states visited during the exploration phase of the task. Line and shading indicate mean and standard error across participants. The increase in thinking time with states visited suggests that humans may also transition to more model based behavior with increasing confidence in the goal location.

**Figure S10:**
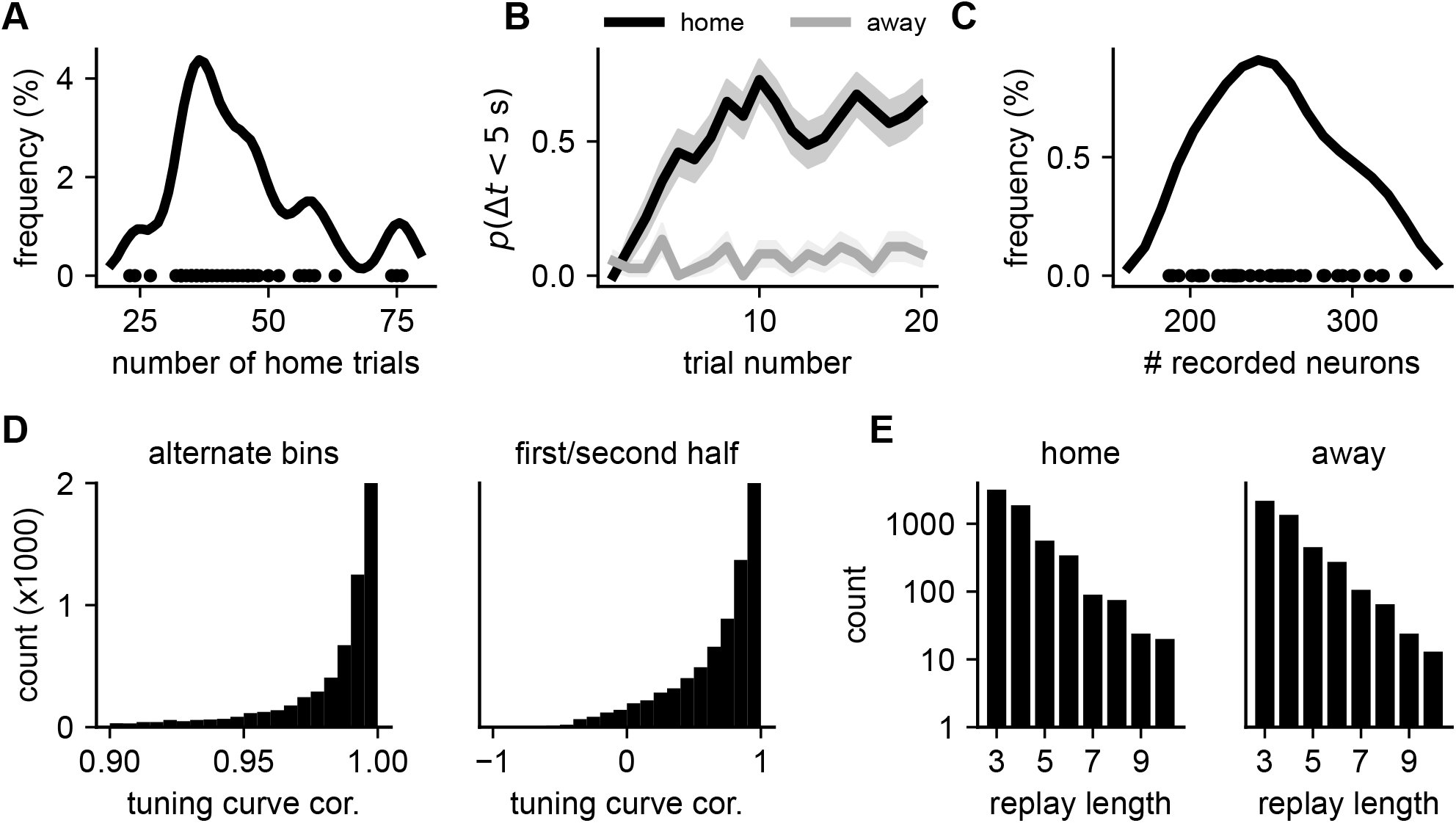
Overview of rodent data. **(A)** Kernel density estimate (*σ* = 3 trials) of the distribution of the number of ‘home’ trials in each session across all animals (an equivalent number of away trials was performed between the home trials). Dots indicate individual sessions. **(B)** Fraction of trials where the animal reached the correct goal location and started licking within 5 seconds of the trial starting, separated by home and away trials. Reaching the goal within 5 seconds was used as a success criterion by Widloski and Foster (2022) since the goal is never explicitly cued at this time (Methods). Line and shading indicate mean and standard error across sessions. The animals learn the location of the home well within a few trials and consistently return to this location on the home trials. **(C)** Distribution of the number of recorded neurons in each session. Line indicates a convolution with a Gaussian filter (15 neuron std) and dots indicate individual sessions. **(D)** Consistency of spatial tuning curves of hippocampal neurons. Consistency was quantified by constructing two tuning curves on the 5 *×*5 spatial grid (Figure 4A) for each neuron and computing the Pearson correlation between the two tuning curves. The data was split into either even/odd time bins in a session (left plot) or first/second half of the session (right plot) to compute a pair of tuning curves. **(E)** Distribution of replay lengths, measured as the number of states visited in a replay, for all replays during home (left) or away (right) trials. Note the log scale on the y-axis.

**Figure S11:**
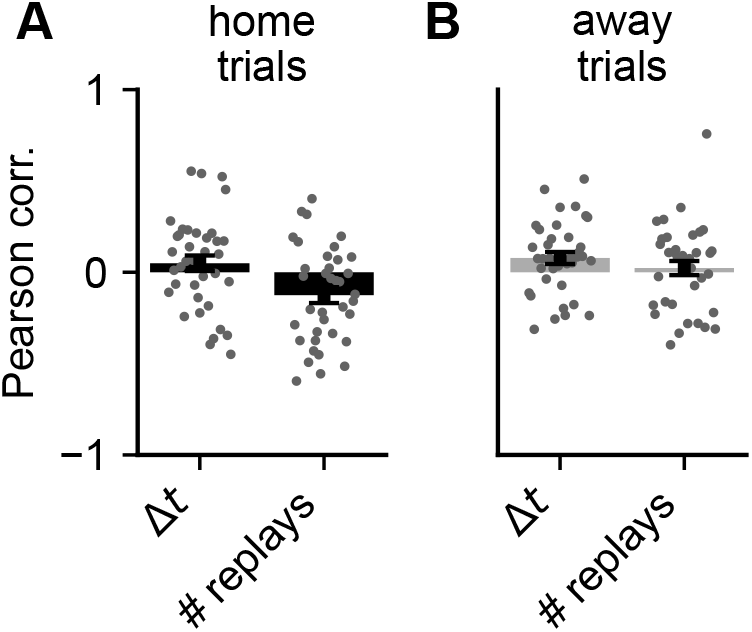
Correlation between distance-to-goal and response time or number of replays in rodents. For the rodent data recorded by Widloski and Foster (2022), we quantified the shortest initial distance-to-goal on each trial as well as (i) the time spent at the previous well before initiating the trial, and (ii) the number of replays recorded during this period. **(A)** Pearson correlation during home trials between initial distance-to-goal and either (i) time spent at the previous well (Δ*t*, left), or (ii) the number of replays performed at the previous well (right). The absence of a correlation between goal distance and time spent at the previous well differs from our analyses of human behavior in a similar maze task (Figure 2C). However, there are two notable differences between these two paradigms that might explain the apparently discrepant results. Firstly, the rats recorded by Widloski and Foster (2022) have to physically consume the reward at the previous well before they can continue their behavior. Secondly, there is an experimenter imposed delay between the end of reward consumption and the next reward becoming available. This is different from the human task, which was explicitly designed to encourage a trade-off between the time spent thinking and the time spent acting, without any additional ‘down time’ that could be used for planning without incurring a temporal opportunity cost. **(B)** As in (A), now for away trials.

**Figure S12:**
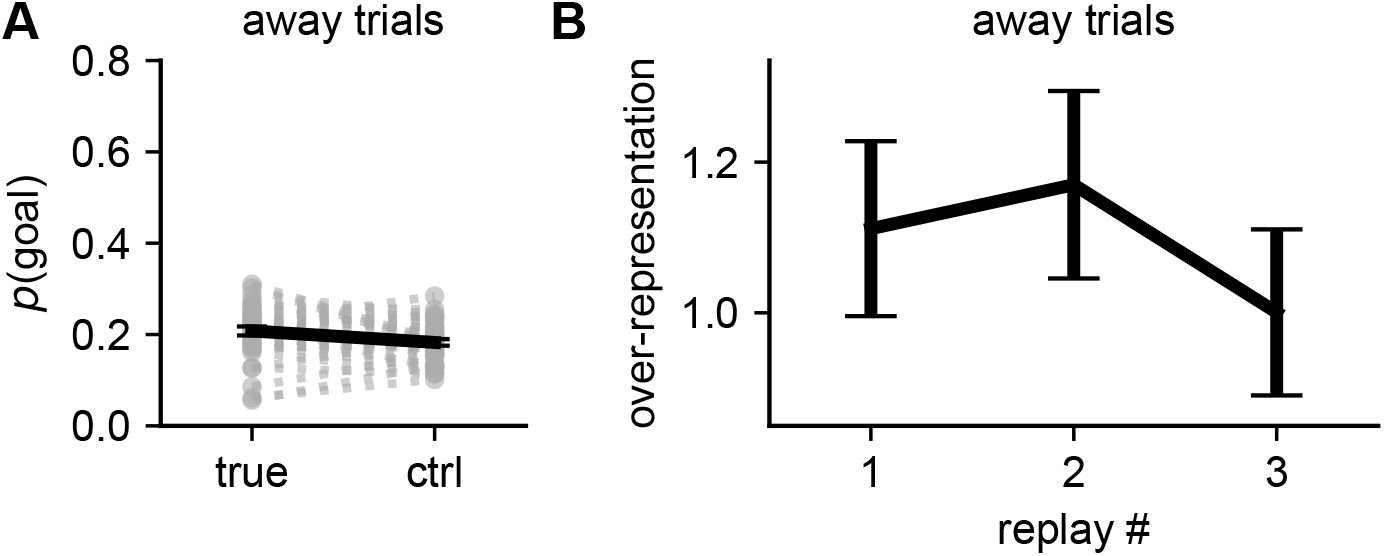
Analyses of replays during away trials. **(A)** Fraction of replays reaching either the true goal (left) or a randomly sampled alternative goal location (right) during away trials. Dashed lines indicate individual sessions (*n* = 37), and solid lines indicate mean and standard error across sessions. In contrast to the home trials (Figure 4C), the goal is not over-represented during away trials, where the goal location is unknown. **(B)** Over-representation of replay success as a function of replay number within sequences of replays containing at least 3 distinct replay events (c.f. Figure 4E). Error bars indicate standard error across replays pooled from all animals. In contrast to the home trials, there is no increase in over-representation with replay number during these away trials.

**Figure S13:**
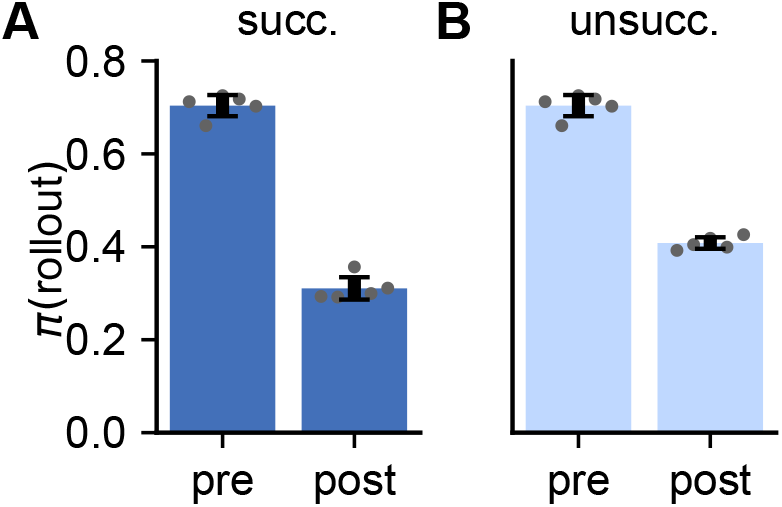
Change in *π*(rollout) for successful and unsuccessful rollouts. **(A)** *π*(rollout) before (left) and after (right) successful rollouts. Bars and error bars indicate mean and standard error across five RL agents (gray dots). The data used for this analysis was the same data used in Figure 3E. **(B)** As in (A), now for unsuccessful rollouts. *π*^post^(rollout) was substantially larger after unsuccessful than successful rollouts (Δ*π*^post^(rollout) = 0.10 *±* 0.01 mean *±* sem).

## Methods

### Software

All models were trained in Julia version 1.7 using Flux and Zygote for automatic differentiation (Innes et al., 2018). Human behavioral experiments were written in OCaml, with the front-end transpiled to javascript for running in the participants’ browsers. All analyses of the models and human data were performed in Julia version 1.8. All analyses of hippocampal replay data were performed in Python 3.8.

### Statistics

Unless otherwise stated, all plots are reported as mean and standard error across human participants (*n* = 94), independently trained RL agents (*n* = 5), or experimental sessions in rodents (*n* = 37).

### Environment

We generated mazes using the following algorithm:

#### Algorithm 1: Maze generating algorithm

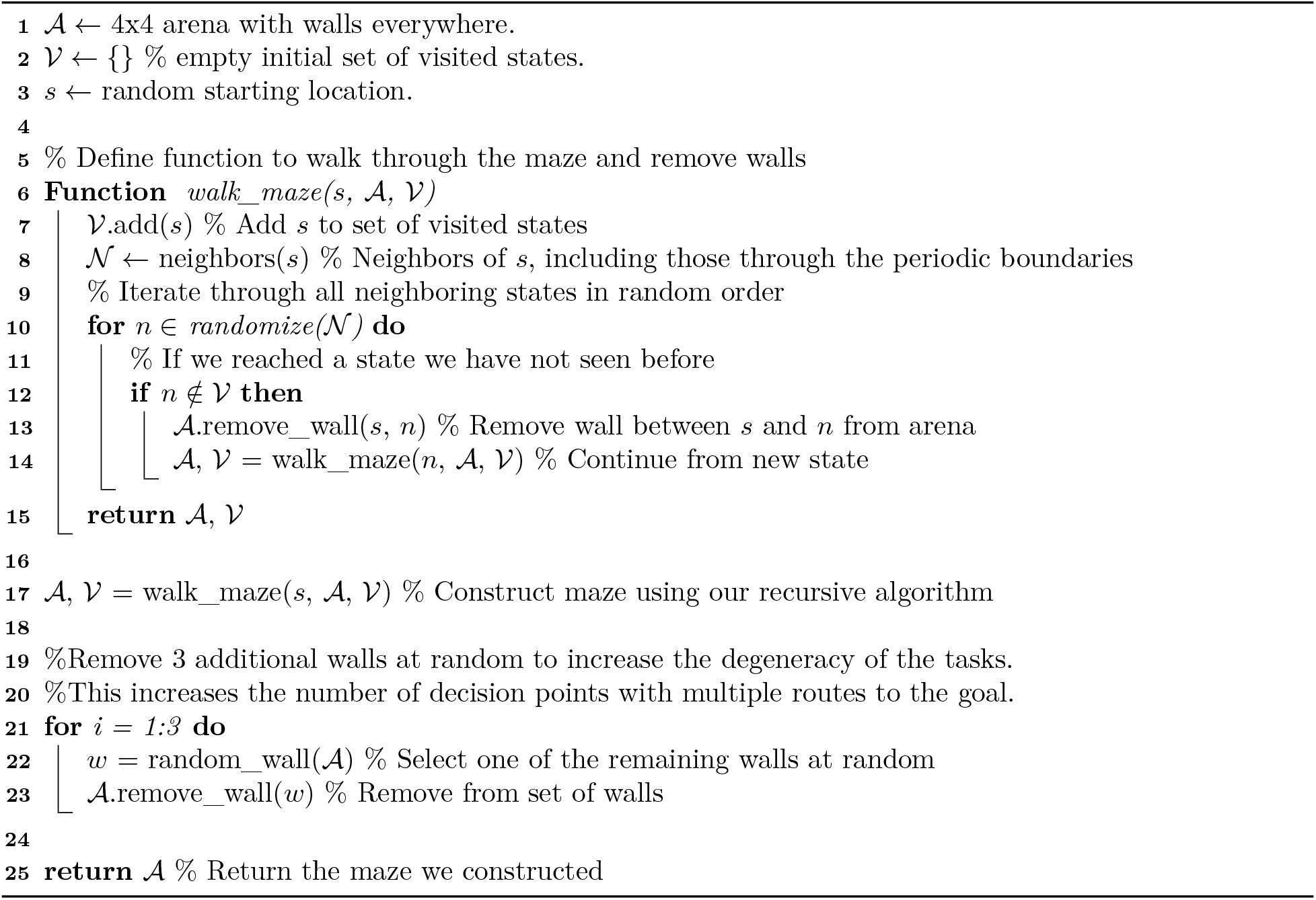

Mazes without periodic boundaries (Figure S1) were generated in the same way, except that states were not considered neighbors across a boundary, and 4 walls were removed instead of 3 walls in the last step of the algorithm in order to approximately match the distributions of shortest paths between pairs of states (Figure S1D).

For each environment, a goal location was sampled uniformly at random. When subjects took an action leading to the goal, they transitioned to this location before being teleported to a random location. In the computational model, this was achieved by feeding the agent an input at this location before teleporting the agent to the new location. The policy of the agent at this iteration of the network dynamics was ignored, since the agent was teleported rather than taking an action.

### Reinforcement learning model

We trained our agent to maximize the expected reward, with the expectation taken both over environments ℰ and the agent’s policy *π*:

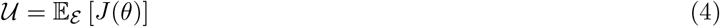

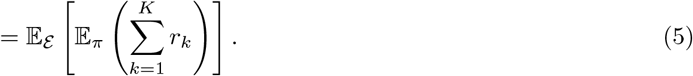

Here, 𝒰 is the *utility function, k* indicates the iteration within an episode, and *r*_*k*_ indicates the instantaneous reward at each iteration. We additionally introduced the following auxiliary losses:

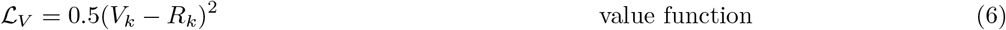

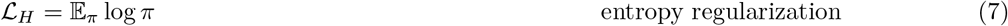

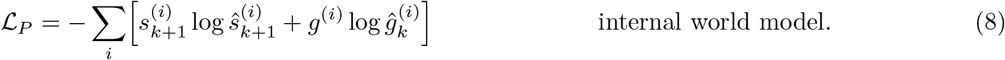

Here, *ĝ*_*k*_, and *ŝ*_*k*+1_ are additional network outputs containing the agent’s estimate of the current reward location and upcoming state as categorical distributions. *g* and *s*_*k*+1_ are the corresponding ground truth quantities, represented as one-hot vectors. 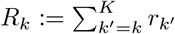 is the empirical cumulative future reward from *k* iteration onwards, and *V*_*k*_ is the value function of the agent.

To maximize the utility and minimize the losses, we trained the RL agent on-policy using a policy gradient algorithm with a baseline (Sutton and Barto, 2018; Jensen, 2023b) and parameter updates of the form

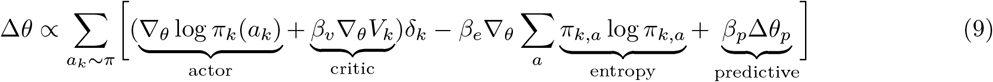

Here, *δ*_*k*_ := *−V*_*k*_ + *R*_*k*_ is the ‘advantage function’, and Δ*θ*_*p*_ = *∇*_*θ*_ ℒ_*P*_ is the derivative of the predictive loss ℒ_*P*_, which was used to train the ‘internal model’ of the agent. *β*_*p*_ = 0.5, *β*_*v*_ = 0.05 and *β*_*e*_ = 0.05 are hyperparameters controlling the importance of the three auxiliary losses. While we use the predictive model explicitly in the planning loop, similar auxiliary losses are also commonly used to speed up training by encouraging the learning of useful representations (Jaderberg et al., 2016).

Our model consisted of a GRU network with 100 hidden units (Cho et al., 2014; Supplementary Note). The policy was computed as a linear function of the hidden state followed by a softmax normalization. The value function was computed as a linear function of the hidden state. The predictions of the next state and reward location were computed with a neural network that received as input a concatenation of the current hidden state ***h***_*k*_ and the action *a*_*k*_ sampled from the policy (as a one-hot representation). The output layer of this feedforward network was split into a part that encoded a distribution over the predicted next state (a vector of 16 grid locations with softmax normalization), and a part that encoded the predicted reward location in the same way. This network had a single hidden layer with 33 units and a ReLU nonlinearity.

The model was trained using ADAM (Kingma and Ba, 2015) on 200,000 batches, each consisting of 40 episodes, for a total of 8 *×*10^6^ training episodes. These episodes were sampled independently from a total task space of (273 *±*13) *×*10^6^ tasks (mean*±* standard error). The total task space was estimated by sampling 50,000 wall configurations and computing the fraction of the resulting 1.25 *×*10^9^ pairwise comparisons that were identical, divided by 16 to account for the possible reward locations. This process was repeated 10 times to estimate a mean and confidence interval. These considerations suggest that the task coverage during training was *≈*2.9%, which confirms that the majority of tasks seen at test time are novel (although we do not enforce this explicitly).

For all evaluations of the model, actions were sampled greedily rather than on-policy unless otherwise stated. This was done since the primary motivation for using a stochastic policy is to explore the space of policies to improve learning, and performance was better under the greedy policy at test time.

### Planning

Our implementation of ‘planning’ in the form of policy rollouts is described in Algorithm 2. This routine was invoked whenever a ‘rollout’ was sampled from the policy instead of a physical action.

#### Algorithm 2: Planning routine for the RL agent

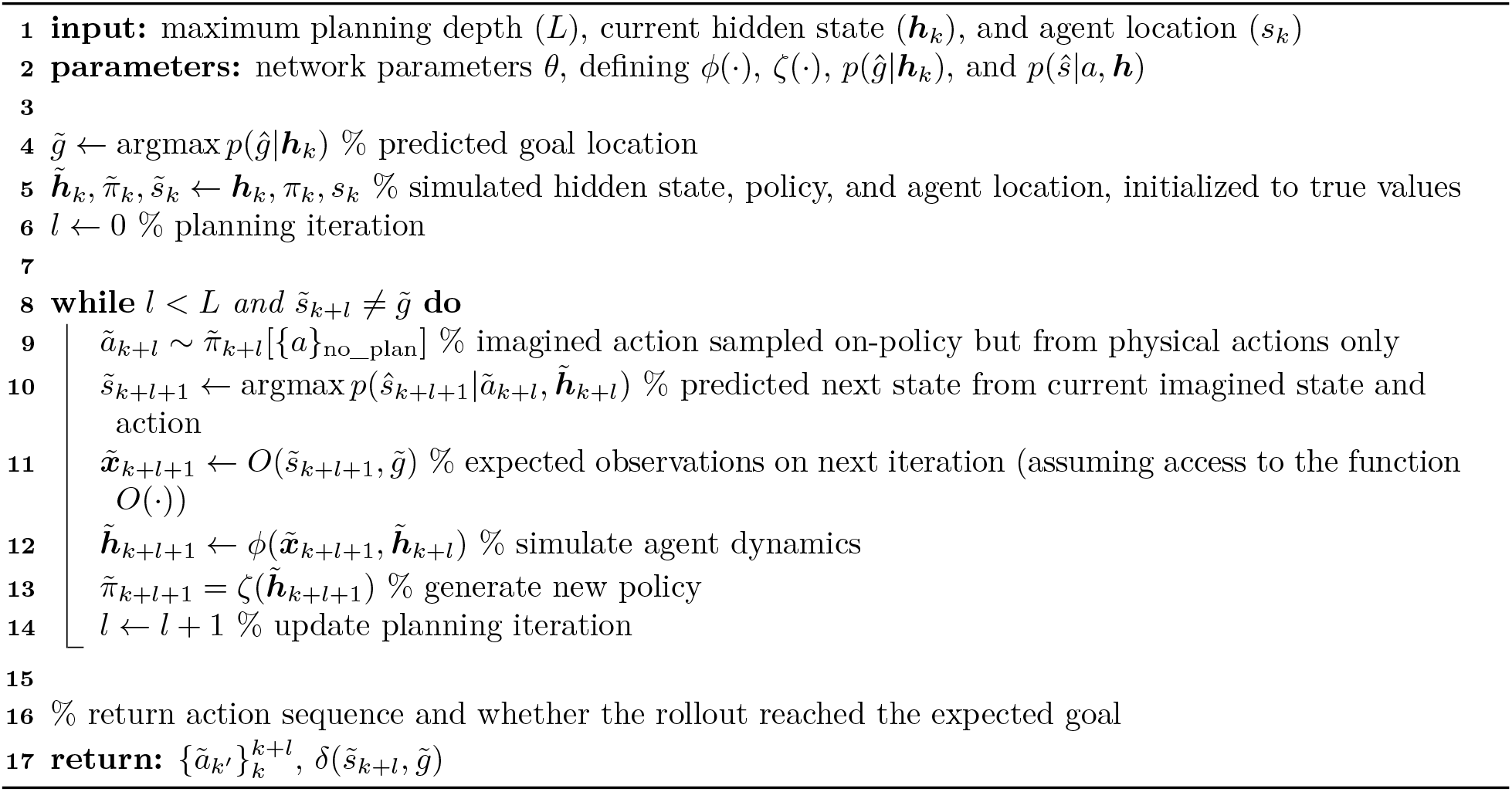

For the network update following a rollout, the input ***x***_*k*+1_ was augmented with an additional ‘rollout input’ consisting of (i) the sequence of simulated actions, each as a 1-hot vector, and (ii) a binary input indicating whether the imagined sequence of states reached the imagined goal location. Additionally, the time within the session was only updated by 120 ms after a rollout in contrast to the 400 ms update after a physical action or teleportation step. For the analyses with a variable temporal opportunity cost of rollouts (Figure S8), we incremented time by *l·* 24 ms after a rollout, where *l* is the number of simulated actions. In Algorithm 2, we assume access to a function 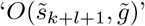, which returns imagined inputs 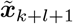. This function is the same as that used to generate inputs from the environment, which means that we assume the ‘predicted’ input to the RNN during a rollout takes the same form as the ‘sensory’ input from the real environment following an action.

While both an imagined ‘physical state’ 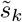 and ‘hidden state’ 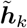 are updated during the rollout, the agent continues from the *original* location *s*_*k*_ and hidden state ***h***_*k*_ after the rollout, but with an augmented input. Additionally, gradients were not propagated through the rollout process, which was considered part of the ‘environment’. This means that there was no explicit gradient signal that encouraged the policy to drive useful or informative rollouts. Instead, the rollout process simply relied on the utility of the base policy optimized for acting in the environment.

### Performance by number of rollouts

To quantify the performance as a function of the number of planning steps in the RL agent (Figure 3A), we simulated each agent in 1,000 different mazes until it first found the goal and was teleported to a random location. We then proceeded to enforce a particular number of rollouts before the agent was released in trial 2. During this release phase, no more rollouts were allowed – in other words, the policy was re-normalized over the physical actions, and the probability of performing a rollout was set to zero. Performance was then quantified as the average number of steps needed to reach the goal during this test phase. For the control without feedback, we repeated this analysis with all feedback from the rollouts set to zero, while the recurrent dynamics were allowed to proceed as usual. The optimal reference value was computed as the average optimal path length for the trial 2 starting states.

When performing more than one sequential rollout prior to taking an action, the policy of the agent can continue to change through two potential mechanisms. The first is that the agent can explicitly ‘remember’ the action sequences from multiple rollouts and somehow arbitrate between them. The second is to progressively update the hidden state in a way that leads to a better expected policy with each rollout, since the feedback from a rollout is incorporated into the hidden state that induces the policy used to draw the next rollout. On the basis of the analysis in Figure 5, we expect the second mechanism to be dominant, although we did not explicitly test the ability of the agent to ‘remember’ multiple action sequences from sequential rollouts. For these and all other RNN analyses, the agent executed the most likely action under the policy during ‘testing’ in contrast to the sampling performed during training, where such stochasticity is necessary for exploring the space of possible actions. All results were qualitatively similar if actions were sampled during the test phase, although average performance was slightly worse.

### Performance in the absence of rollouts and with shuffled rollout times

To quantify the performance of the RL agent in the absence of rollouts, we let the agent receive inputs and produce outputs as normal. However, we set the probability of performing a rollout under the policy to zero and re-normalized the policy over the physical actions before choosing an action from the policy. We compared the average performance of the agent (number of rewards collected) in this setting to the performance of the default agent in the same environments.

To compare the original performance to an agent with randomized rollout times, we counted the number of rollouts performed by the default agent in each environment. We then resampled a new set of network iterations at which to perform rollouts, matching the size of this new set to the original number of rollouts performed in the corresponding environment. Finally, we let the agent interact with the environment again, while enforcing a rollout on these network iterations, and preventing rollouts at all other timesteps. It is worth noting that we could not predict *a priori* the iterations on which the agent would find the goal, at which point rollouts were not possible. If a rollout had been sampled at such an iteration, we resampled this rollout from the set of remaining network iterations.

### Rollouts by network size

To investigate how the frequency of rollouts depended on network size (Figure 5E; Figure S3), we trained networks with either 60, 80, or 100 hidden units (GRUs). Five networks were trained of each size. At regular intervals during training, we tested the networks on a series of 5,000 mazes and computed (i) the average reward per episode, and (ii) the fraction of actions that were rollouts rather than physical actions. We then plotted the rollout fraction as a function of average reward to see how frequently an agent of a given size performed rollouts for a particular performance.

### Effect of rollouts on agent policy

To quantify the effect of rollouts on the policy of the agent, we simulated each agent in 1,000 different mazes until it first found the goal and was teleported to a random location. We then resampled rollouts until both (i) a successful rollout and (ii) an unsuccessful rollout had been sampled. Finally, we quantified *π*^pre^(*â*_1_) and *π*^post^(*â*_1_) separately for the two scenarios and plotted the results in Figure 3E. Importantly, this means that each data point in the ‘successful’ analysis had a corresponding data point in the ‘unsuccessful’ analysis with the exact same maze, location, and hidden state. In this way, we could query the effect of rollouts on the policy without the confound of how the policy itself affects the rollouts. For this analysis, we discarded episodes where the first 100 sampled rollouts did not result in both a successful and an unsuccessful rollout.

For Figure S13, we used the same episodes and instead quantified *π*(rollout) before and after the rollout, repeating the analysis for both successful and unsuccessful rollouts.

### Overlap between hidden state updates and policy gradients

Using a single rollout 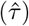 to approximate the expectation over trajectories of the gradient of the expected future reward for a given episode, *∇*_***h***_*J*_fut_(***h***), the policy gradient update in ***h*** takes the form 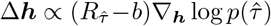. Here, Δ***h*** is the change in hidden state resulting from the rollout, 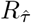 is the ‘reward’ of the simulated trajectory, *b* is a constant or state-dependent baseline, and 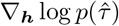 is the gradient with respect to the hidden state of the log probability of 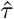 under the policy induced by ***h***. This implies that the *derivative* of the hidden state update w.r.t. 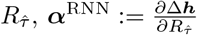,should be proportional to 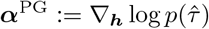.

For these analyses, we divided 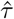 into its constituent actions, defining 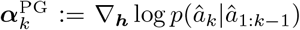 as the derivative w.r.t. the hidden state of the log probability of taking the simulated action at step *k*, conditioned on the actions at all preceding steps (1 to *k −*1) being consistent with the rollout. To compute ***α***^RNN^, we also needed to take derivatives w.r.t. 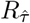 – the ‘reward’ of a rollout. A naive choice here would be to simply consider 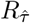 to be the input specifying whether the rollout reached the reward or not. However, we hypothesized that the agent would also use information about e.g. how long the simulated trajectory was in its estimate of the ‘goodness’ of a rollout (since a shorter rollout implies that the goal was found faster). We therefore determined the direction in planning input state space that was most predictive of the time-to-goal of the agent. We did this by using linear regression to predict the (negative) time-to-next-reward as a function of the planning feedback ***x***_*f*_ across episodes and rollouts. This defines the (normalized) direction 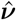 in planning input space that maximally increases the expected future reward. Finally, we defined 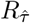 as the magnitude of the planning input in direction 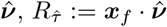. We could then compute ***α***^RNN^ with this definition of 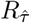 using automatic differentiation.

In Figure 5C, we computed ***α***^RNN^ and 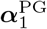 across 1,000 episodes and subtracted the mean across rollouts for each feature. We then performed PCA on the set of 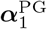 and projected both ***α***^RNN^ and 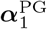 into the space spanned by the top 3 PCs. Finally, we computed the mean value of both quantities conditioned on *â*_1_ to visualize the alignment. In Figure 5D, we considered the same ***α***^RNN^ and 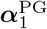 vectors. After mean subtraction for each feature and normalization across features for each ***α***, we projected these into the space spanned by the top 3 PCs of 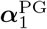. Finally, we computed the average across rollouts of the cosine similarity between the pairs of 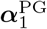 and 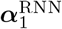 in this latent space. We performed this analysis in a low-dimensional space because we were primarily interested in changes to ***h*** within the subspace that would affect 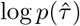. As a control, we repeated the analysis after altering the planning input ***x***_*f*_ to falsely inform the agent that it had simulated some other action *â*_1,ctrl_ ≠ *â*_1_. Finally, we also repeated this analysis using 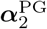 to characterize how the effects of the planning input propagated through the recurrent network dynamics to modulate future action probabilities.

### Quantification of value functions

To quantify the error of the value function in Figure S6, we compared the value function computed by the agent (*V*_*k*_) to the true reward-to-go 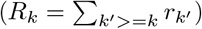. Figure S6A shows the distribution of errors *V*_*k*_*− R*_*k*_, while the ‘constant control’ shows the distribution of 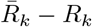, where 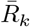 is the mean reward-to-go across all trials and iterations. These distributions were aggregated across all agents. In Figure S6C, we considered sequences of *N* consecutive rollouts and computed the average value function before the first rollout and after each rollout. Figure S6D further conditions on all the rollouts in a sequence being ‘unsuccessful’.

### Human data collection

The human behavioral experiments used in this study have been certified as exempt from IRB review by the UC San Diego Human Research Protection Program. We collected data from 100 human participants (50 male, 50 female) recruited on Prolific to perform the task described in Figure 1B. All participants provided informed consent prior to commencing the experiment. Subjects were asked to complete (i) 6 ‘guided’ episodes where the optimal path was shown explicitly, followed by (ii) 40 non-guided episodes, and (iii) 12 guided episodes. The task can be found online. During data collection, a subject was deemed ‘disengaged’, and the trial repeated, if one of three conditions were met: (i) the same key was pressed 5 times in a row, (ii) the same key pair was pressed four times in a row, or (iii) no key was pressed for 7 seconds. Participants were paid a fixed rate of $3 plus a performance-dependent bonus of $0.002 for each completed trial across both guided and non-guided episodes. The experiment took approximately 22 minutes to complete, and the average pay across participants was $10.5 per hour including the performance bonus. For the experiment without periodic boundaries (Figure S1), we collected data from 49 human participants (25 male, 24 female). The experiment was performed as described above, with the only difference being that we used mazes where participants could not move through the boundaries.

The data from 6 participants with a mean response time greater than 690 ms during the guided episodes were excluded to avoid including participants who were not sufficiently engaged with the task. For the guided episodes, only the last 10 episodes were used for further analyses. For the non-guided episodes, we discarded the first two episodes and used the last 38 episodes. This was done to give participants two episodes to get used to the task for each of the two conditions, and the first set of guided episodes was intended as an instruction in how to perform the task.

### Performance as a function of trial number

We considered all episodes where the humans or RL agents completed at least four trials, evaluating the RL agents across 50,000 episodes. We then computed the average across these episodes of the number of steps to goal as a function of trial number separately for all subjects. Figure 2A illustrates the mean and standard error across subjects (human participants or RL agents). The optimal value during the exploitation phase was computed by using dynamic programming to find the shortest path between each possible starting location and the goal location, averaged across all environments seen by the RL agent. To compute the exploration baseline, brute force search was used to identify the path that explored the full environment as fast as possible. The optimal exploration performance was then computed as the expected time-to-first-reward under this policy, averaged over all possible goal locations.

### Estimation of thinking times

In broad strokes, we assumed that for each action, the response time *t*_r_ is the sum of a thinking time *t*_t_ and some perception-action delay *t*_d_, both subject to independent variability:

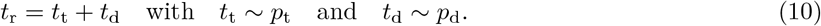

Here, *{t*_r_, *t*_t_, *t*_d_*}≥* 0 since elapsed time cannot be negative. We assumed that the prior distribution over perception-action delays, *p*_d_, was identical during guided and non-guided trials. For each subject, we obtained a good model of *p*_d_ (see below) by considering the distribution of response times measured during guided trials. This was possible because guided trials involved no ‘thinking’ by definition, such that *t*_d_ *≡ t*_r_ was directly observed. Finally, for any non-guided trial with observed response *t*_r_, we formed a point estimate of the thinking time by computing the mean of the posterior *p*(*t*_t_|*t*_r_):

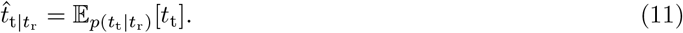

In more detail, we took *p*_t_ during non-guided trials to be uniform between 0 and 7 s – the maximum response time allowed, beyond which subjects were considered disengaged, and the trial was discarded and reset. For *p*_d_(*t*_d_), we assumed a shifted log-normal distribution,

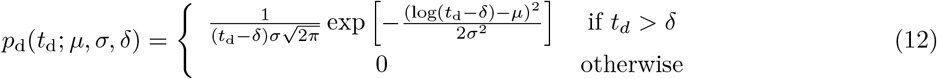

with parameters *μ, σ*, and *δ* obtained via maximum likelihood estimation based on the collection of response times *t*_r_ *≡ t*_d_ observed during guided trials. For a given *δ*, the maximum likelihood values of *μ* and *σ* are simply given by the mean and standard deviation of the logarithm of the shifted observations. To fit this shifted log-normal model, we thus performed a grid search over 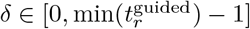 at 1 ms resolution and selected the value under which the optimal (*μ, σ*) gave the largest likelihood. This range of *δ* was chosen to ensure that (i) only positive values of 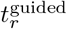 had positive probability, and (ii) all observed 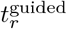 had non-zero probability. We then retained the optimal *μ, σ*, and *δ* to define the prior over *p*_d_(*t*_*d*_) on non-guided trials for each subject.

According to Bayes’ rule, the posterior is proportional to

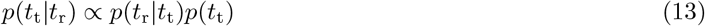

where

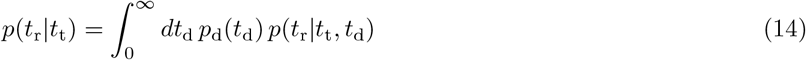

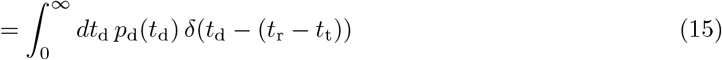

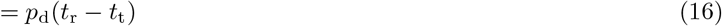

Therefore, the posterior is given by

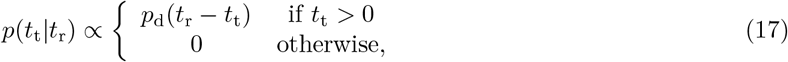

resulting in the following posterior mean:

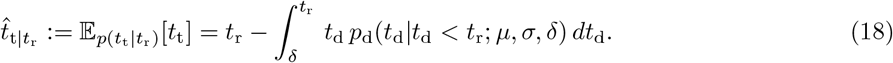

Here, *p*_d_(*t*_d_ | *t*_d_ *< t*_r_) denotes *p*_d_(*t*_d_) re-normalized over the interval *t*_d_ *< t*_r_, and the condition (*t*_d_ *< t*_r_) is equivalent to (*t*_t_ *>* 0). We note that the integral runs from *δ* to *t*_r_ since *p*_d_(*t*_d_) = 0 for *t*_d_ *< δ*. As *δ* simply shifts the distribution over *t*_d_, we can rewrite this as

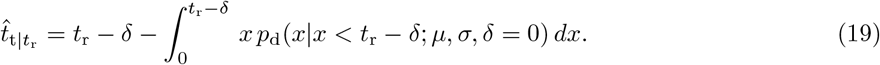

This is useful since the conditional expectation of a log-normally distributed random variable with *δ* = 0 is given in closed form by

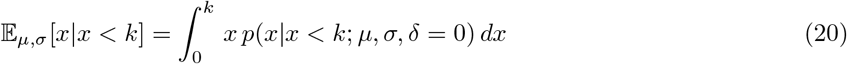

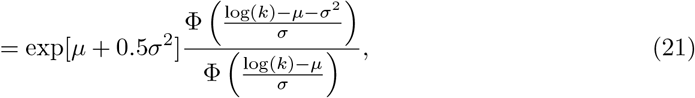

where Φ(*·*) is the cumulative density function of the standard Gaussian, 𝒩(0, 1). This allows us to compute the posterior mean thinking time for an observed response time *t*_r_ in closed form as

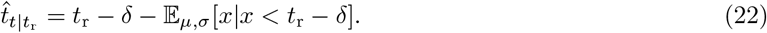

We note that the support of *p*_d_(*t*_d_| *t*_d_ *< t*_r_; *μ, σ, δ*) is *t*_d_ *∈* [*δ, t*_r_]. For 0.6% of the non-guided decisions, the value of *t*_r_ was *lower* than the estimated *δ* for the corresponding participant, in which case *p*(*t*_t_ |*t*_r_) is undefined. In such cases, we defined the thinking time to be 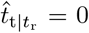, since the response time was shorter than our estimated minimum perception-action delay. A necessary (but not sufficient) condition for *t*_r_ *< δ* is that *t*_r_ is smaller than the smallest response time in the guided trials.

The whole procedure of fitting and inference described above was repeated separately for actions that immediately followed a teleportation step (i.e. the first action in each trial) and for all other actions. This is because we expected the first action in each trial to be associated with an additional perceptual delay compared to actions that followed a predictable transition.

While this approach dissociates thinking from other forms of sensorimotor processing to some extent, the ‘thinking times’ reported in this work still only represent a best estimate given the available data, and we use ‘thinking’ to refer to any internal computational process guiding decision making. This does not necessarily imply a conscious process that we can introspect, since decision making occurs on a fast timescale of hundreds of milliseconds.

All results were qualitatively similar using other methods for estimating thinking time, including (i) a log-normal prior over *t*_d_ with no shift (*δ* = 0), (ii) using the posterior mode instead of the posterior mean, (iii) estimating a constant *t*_d_ from the guided trials, and (iv) estimating a constant *t*_d_ as the 0.1 or 0.25 quantile of *t*_r_ from the non-guided trials.

### Thinking times in different situations

To investigate how the thinking time varied in different situations, we considered only exploitation trials and computed for every action (i) the minimum distance to the goal *at the beginning of the corresponding trial*, and (ii) what action number this was within the trial. We then computed the mean thinking time as a function of action number separately for each initial distance-to-goal. This analysis was repeated across experimental subjects and results reported as mean and standard error across subjects.

We repeated this analysis for the RL agents, where ‘thinking time’ was now defined based on the average number of rollouts performed, conditioned on action-within-trial and initial distance to goal.

### Comparison of human and model thinking times

For each subject and each RL agent, we clamped the trajectory of the agent to that taken by the subject (i.e. we used the human actions instead of sampling from the policy). After taking an action, we recorded *π*(rollout) under the model on the first timestep of the new state for comparison with human thinking times. We then sampled a rollout with probability *π*(rollout) and took an action (identical to the next human action) with probability 1 *− π*(rollout), repeating this process until the next state was reached. Finally, we computed the average *π*(rollout) across 20 iterations of each RL agent for comparison with the human thinking time in each state. Figure 2E shows the human thinking time as a function of *π*(rollout), with the bars and error bars illustrating the mean and standard error in each bin. For this analysis, data was aggregated across all participants. Results were similar if we compared human thinking times with the average number of rollouts performed rather than the initial *π*(rollout).

In Figure 2F, we computed the correlation between thinking time and various regressors on a participant-by-participant basis and report the result as mean and standard error across participants (*n* = 94). For the ‘residual’ correlation, we first computed the mean thinking time for each momentary distance-to-goal for each participant and the corresponding mean *π*(rollout) for the RL agents. We then subtracted the appropriate mean values from the thinking times and *π*(rollout) in the human participants and RL agents. In other words, we subtracted the average thinking time across all situations 5 steps from the goal from all individual data points where the participant was 5 steps from the goal etc. Finally, we computed the correlation between the residual *π*(rollout) and the residual thinking times. This analysis was repeated across all participants and the result reported as mean and standard error across participants. Note that all ‘distance-to-goal’ measures refer to the shortest path to goal rather than the number of steps actually taken by the participant to reach the goal.

### Analysis of hippocampal replays

For our analyses of hippocampal replays in rats, we used data recently recorded by Widloski and Foster (2022). This dataset consisted of a total of 37 sessions from 3 rats (n = 17, 12, 8 sessions for each rat) as they performed a dynamic maze task. This task was carried out in a square arena with 9 putative reward locations. In each session, six walls were placed in the arena, and a single reward location was randomly selected as the ‘home’ well. The task involved alternating between moving to this home well and a randomly selected ‘away’ well. Importantly, a delay of 5-15 s was imposed between the animal leaving the previous rewarded well before reward (chocolate milk) became available at the next rewarded well. On the away trials, the emergence of reward was also accompanied by a visual cue at the rewarded well, informing the animal that this was the reward location. We only considered replays at the previous well before this visual cue and reward became available. In a given session, the animals generally performed around 80 trials (40 home trials and 40 away trials; Figure S10). For further task details, we refer to Widloski and Foster (2022).

For our analyses, we only included trials that lasted less than 40 seconds. We did this to discard time periods where the animals were not engaged with the task. Additionally, we discarded the first home trial of each session, where the home location was unknown, since we wanted to compare the hippocampal replays with model rollouts during the exploitation phase of the maze task. For all analyses, we discretized the environment into a 5*×* 5 grid (the 3*×* 3 grid of wells and an additional square of states around these) in order to facilitate more direct comparisons with our human and RNN task. Following Widloski and Foster (2022), we defined ‘movement epochs’ as times where the animal had a velocity greater than 2 cm/s and ‘stationary epochs’ as times there the animal had a velocity less than 2 cm/s.

### Replay detection

To detect replays, we followed Widloski and Foster (2022) and fitted a Bayesian decoder to neural activity as a function of position during movement epochs in each session, assuming Poisson noise statistics and considering only neurons with an average firing rate of at least 0.1 Hz over the course of the session. This decoder was trained on a rolling window of neural activity spanning 75 ms and sampled at 5 ms intervals (Widloski and Foster, 2022). We then detected replays during stationary epochs by classifying each momentary hippocampal state as the maximum likelihood state under the Bayesian decoder, again using neural activity in 75 ms windows at 5 ms intervals. Forward replays were defined as sequences of states which included 2 consecutive transitions to an adjacent state (i.e. a temporally and spatially contiguous sequence of three or more states), and which originated at the true animal location. For all animals, we only analyzed replays where the animal was at the previous reward location *before* it initiated the new trial (c.f. Widloski and Foster, 2022). To increase noise robustness, we allowed for short ‘lapses’ in a replay, defined as periods with a duration less than or equal to 20 ms, where the decoded location moved to a distant location before returning to the previously decoded location. These lapses were ignored for downstream analyses.

### Wall avoidance

To compute the wall avoidance of replays (Figure 4B), we calculated the fraction of state transitions that passed through a wall. This was done across all replays preceding a ‘home’ trial (i.e. when the animal knew the next goal). As a control, we computed the same quantity averaged over 7 control conditions, which corresponded to the remaining non-identical rotations and reflections of the walls from the corresponding session. We repeated this analysis for all sessions and report the results in Figure 4 as mean and standard error across sessions. To test for significance, we randomly permuted the ‘true’ and ‘control’ labels independently for each session and computed the fraction of permutations (out of 10,000), where the difference between ‘control’ and ‘true’ was larger than the experimentally observed value. This analysis was also repeated in the RL agent, where the control value was computed with respect to 50,000 other wall configurations sampled from the maze generating algorithm (Algorithm 1).

### Reward enrichment

To compute the reward enrichment in hippocampal replays (Figure 4C), we computed the fraction of all replays preceding a ‘home’ trial that passed through the reward location. As a control, we repeated this analysis for the remaining 7 locations that were neither the reward location nor the current agent location (for each replay). Control values are reported as the average across these 7 control locations across all replays. This analysis was repeated for all sessions. While this can lead to systematic differences between true and control values in individual trials depending on how close the true reward location is to the current animal locations, the distance-to-goal will be the same in expectation between the true reward location and the control locations. This is also why we do not see an effect for the away trials in Figure S12A.

To test for significance, we randomly permuted the ‘goal’ and ‘control’ labels independently for each session. Here, the ‘goal’ and ‘control’ values permuted were the ones computed by averaging across all trials and control locations in the session (i.e. we randomly swapped the 37 pairs of data points shown in gray in Figure 4C). We then computed the fraction of permutations (out of 10,000) where the difference between ‘goal’ and ‘control’ was larger than the experimentally observed value after averaging across sessions.

This analysis was also repeated in the RL agent, where the control value was computed across the 14 locations that were not the current agent location or true goal.

### Behavior by replay type

To investigate how the animal behavior depended on the type of replay (Figure 4D), we analyzed home trials and away trials separately. We constructed a list of all the ‘first’ replayed actions *â*_1_, defined as the cardinal direction corresponding to the first state transition in each replay. We then constructed a corresponding list of the first physical action following the replay, corresponding to the cardinal direction of the first physical state transition after the replay. Finally, we computed the overlap between these two vectors to arrive at the probability of ‘following’ a replay. This overlap was computed separately for ‘successful’ and ‘unsuccessful’ replays, where successful replays were defined as those that reached the goal without passing through a wall. For the unsuccessful replays, we considered the 7 remaining locations that were not the current animal location or current goal. We then computed the average overlap under the assumption that each of these locations were the goal, while discarding replays that were successful for the ‘true’ goal. The reason for not considering replays that were successful for the ‘true goal’ in the ‘unsuccessful’ setting is because we are primarily interested in the distinction between replays that are ‘successful’ vs. ‘unsuccessful’ to the true goal and therefore wanted disjoint sets of replays in these two analyses. However, we did this while considering whether replays were ‘successful’ towards a control location in order to better match the spatiotemporal statistics of replays in the two categories. The analysis was performed independently across all sessions and results reported as mean and standard error across sessions. To test for significance, we randomly permuted the ‘successful’ and ‘unsuccessful’ labels independently for each session and computed the fraction of permutations (out of 10,000) where the difference between successful and unsuccessful replays was larger than the experimentally observed value.

To confirm that our results were not biased by the choice to exclude replays that were successful to the true goal location from our set of ‘unsuccessful’ replays, we performed an additional control analysis, where the control replays were the full set of replays that were ‘successful’ towards a randomly sampled control location, concatenated across control locations. In this case, the control value for the home trials was *p*(*a*_1_ = *â*_1_) = 0.433 instead of *p*(*a*_1_ = *â*_1_) = 0.403 with the disjoint set of ‘unsuccessful’ replays used in the main text. This is still significantly smaller than the value of *p*(*a*_1_ = *â*_1_) = 0.622 for the true goal location, with our permutation test in both cases yielding *p <* 0.001 for the home trials and no significant effect for the away trials.

This analysis was also repeated in the RL agent, where we considered all exploitation trials together since they were not divided into ‘home’ or ‘away’ trials. In this case, the control was computed with respect to all 14 locations that were not the current location or current goal location.

### Effect of consecutive replays

To compute how the probability of a replay being ‘successful’ depended on replay number (Figure 4E), we considered all trials where an animal performed at least 3 replays. We then computed a binary vector indicating whether each replay was successful or not. From this vector, we subtracted the expected success frequency from a linear model predicting success from (i) the time since arriving at the current well, and (ii) the time until departing the current well. We did this to account for any effect of time that was separate from the effect of replay number, since such an effect has previously been reported by Ólafsdóttir et al. (2017). However, this work also notes that many of what they denote ‘disengaged’ replays are non-local and would automatically be filtered out by our focus on local replays. When fitting this linear model, we capped all time differences at a maximum value of |Δ*t*| = 15 s to avoid the analysis being dominated by outliers, and because Ólafsdóttir et al. (2017) only observe an effect for time differences in this range. Our results were not sensitive to altering or removing this threshold. We then conditioned on replay number and computed the probability of success (after regressing out time) as a function of replay number. Finally, we repeated this analysis for all 7 control locations for each replay and divided the true values by control values defined as the average across replays of the average across control locations. A separate correction factor was subtracted from these control locations, which was computed by fitting a linear model to predict the average probability of successfully reaching a control location as a function of the predictors described above. The normalization by control locations was done to account for changes in replay statistics that might affect the results, such as systematically increasing or decreasing replay durations with replay number. To compute the statistical significance of the increase in goal over-representation, we also performed this analysis after independently permuting the order of the replays in each trial to break any temporal structure. This permutation was performed *after* regressing out the effect of time. We repeated this analysis across 10,000 independent permutations and computed statistical significance as the number of permutations for which the increase in over-representation was greater than or equal to the experimental value.

For the corresponding analysis in the RL agents, we did not regress out time since there is no separability between time and replay number. Additionally, the RL agent cannot alter its policy in the absence of explicit network updates – which in our model are always tied to either a rollout or an action. As noted in the main text, an increase in the probability of ‘success’ with replay number in the RL agent could also arise from the fact that performing further replays is less likely after a successful replay than after an unsuccessful replay (Figure S13). We therefore performed the analysis of consecutive replays in the RL agent in a ‘cross-validated’ manner at the level of the policy. In other words, every time the agent performed a rollout, we drew two samples from the rollout generation process. The first of these samples was used as normal by the agent to update ***h***_*k*_ and drive future behavior. The second sample was never used by the agent, but was instead used to compute the ‘success frequency’ for our analyses. This was done to break the correlation between the choice of performing a replay and the assessment of how good the policy was, which allowed us to compute an unbiased estimate of the quality of the policy as a function of replay number. As mentioned in the main text, such an analysis is not possible in the biological data. However, since the biological task was not a reaction time task, we expect less of a causal effect of replay success on the number of replays. Additionally, as noted in the text, if some of the effect in the biological data is in fact driven by a decreased propensity for further replays after a successful replay, that is in itself supporting evidence for a theory of replays as a form of planning.

## Discussion of experimental and architectural choices

In this note, we discuss some of the many architectural and modeling choices that went into our work. As is the case for much work in modern computational neuroscience, the space of models was vast – and larger than we could feasibly explore fully in a single paper. In what follows, we hope to provide some additional motivation for the choices that were made in the main paper and to provide additional intuition for the importance and effect of various architectural choices and hyperparameters in our work. This note is also unlikely to be exhaustive, but we hope that it will be useful both for the reader hoping to gain a deeper understanding of our work, and for those looking to draw inspiration from it in their own computational models.

### Network size

The size of the network used in our work is of some importance. We show in Figure S5 that our key results hold across a range of different network sizes. However, as the network becomes larger, its model-free ‘base policy’ also becomes better – to the point where rollouts become less and less useful as there is less room for improvement from the base policy. Indeed, in the limit of an infinitely large network trained on an infinitely large dataset, we expect a perfect base policy and no rollouts. On the contrary, if the network gets too small, it is unlikely to be able to learn how to use the rollouts for policy improvement, and we again expect rollouts to be less useful. In both limits, we also expect the notions of ‘large’ and ‘small’ to depend on the complexity of the task in question. While it may seem like a limitation of this work that the results ‘only’ hold for certain choices of task and network size, we believe that this is consistent with how humans and other animals operate. In particular, if we are given a particularly easy task, like navigation to a location 100 meters down the road, we are unlikely to spend any time thinking. Similarly, if we are given a seemingly impossible task, like solving a very long and complicated equation, we may simply make a guess without taking the time to think through every step. These notions of ‘very easy’ and ‘very difficult’ are also likely to differ across animals with different cognitive capabilities.

For the task considered in this study, we found substantial use of rollouts across a range of network sizes between 40 to 140, and we found that the frequency of rollouts tended to decrease with network size (Figure S3). We did not test any networks larger than 140 units due to computational constraints associated with the training of large networks. Networks smaller than 40 units also exhibited reduced rollout frequencies, which we hypothesize is because they do not have the capacity to learn both how to solve the task and how to take advantage of the rollout machinery. The exact range of sizes for which our results hold will also depend on the type of network used, with LSTMs likely to be similar to the GRUs used in this work, and vanilla RNNs probably needing larger networks for comparable performance.

### Planning horizon

In this work, we assumed a constant maximum planning horizon of *L* = 8 steps and showed in Figure S5 that our key results are robust to changes to this hyperparameter between 4 and 12 (we did not test values outside this range). We chose a value of 8 for the main paper since it seemed like a reasonable planning depth in our fairly simple task with a relatively small action space, and it is comparable to the planning depth estimated in other simple games (van Opheusden et al., 2023). It is worth noting that some aspects of our results do change with planning depth. In particular, the change in policy for successful and unsuccessful rollouts is dependent on *L*, such that longer planning horizons lead to smaller average policy changes for successful rollouts and larger changes for unsuccessful rollouts. In the language of policy gradients, we expect that this is because the ‘baseline’ implicitly used by the network in its state update is related to the average success of a rollout. In other words, if almost all rollouts are successful (because *L* is sufficiently large), little is learned by observing that a rollout is successful, and the policy should not change much on the basis of this information. It is possible that there will be larger average policy changes in this setting if we instead condition on how late in the rollout the reward was found, which contains more information about the ‘goodness’ of the replayed trajectory. It would also be possible to make the planning horizon *variable* and let the agent itself choose its planning depth. This could be done in two different ways, namely by (i) making the agent decide up front how long a trajectory it wants to simulate, or (ii) letting the agent decide in closed-loop by iteratively returning a partial plan and deciding whether to continue planning or terminate and take a physical action. We opted for the simple fixed length solution since it has a smaller action space and fewer network iterations, making optimization easier. However, a variable planning length model may be closer to human behavior and could be an interesting avenue for future research.

### Time cost of acting and planning

Related to the discussion of planning depth, we also assumed a constant temporal opportunity cost of planning for the agent in the main text (see Figure S8 for results with a variable temporal opportunity cost). This was done despite the rollouts having variable length depending on whether and when the goal was reached during the rollout. We did this because the agent did not know *a priori* how long the rollout would be and had no direct control over its length. In the case of hippocampal replays being contained in sharp-wave ripples (SWRs), this is consistent with an assumption that a single trajectory occurs in a single SWR, and that the inter-SWR interval is independent of the length of the replayed trajectory.

More specifically, we defined a rollout in the model to last 120 ms. This is similar to the duration of human hippocampal replays reported in the literature (Kurth-Nelson et al., 2016). In contrast, a single model action was defined to take 400 ms. These values were not directly fitted to the human data, as all model hyperparameters, including the episode length and relative cost of planning and acting, were chosen before any analyses of human behavior to avoid overfitting. Instead, the relative cost of rollouts compared to actions in the model, *β*_roll_ := Δ*t*_rollout_*/*Δ*t*_action_ = 0.3, was chosen such that there was regular use of rollouts in the task. The episode length *T* = 50 *actions* was chosen to facilitate training of the model. We then designed our human behavioral experiment in a way that allowed participants to take approximately the same number of actions in a given episode as the model, which motivated an episode length of *T* = 20 *seconds*. This implicitly defined the ‘duration’ of a model action as Δ*t*_action_ = 20 seconds*/*50 actions = 400 ms. The duration of a rollout was then defined as Δ*t*_rollout_ = *β*_roll_ *×* 400 ms = 120 ms. Since we did not explicitly fit these parameters to the data, there are likely to be a range of parameter choices that lead to better fits to the human data from this particular experiment. Similarly, there is definitely a range of hyperparameters that lead to worse data fits. Indeed our goal was not to chase the lowest possible discrepancy from human response times, but rather to demonstrate the general concept that models with the ability to perform rollouts do so in similar situations to humans. This is also the reason that we focus on correlations in the paper rather than e.g. MSEs, since the model ‘thinking times’ can be stretched, compressed, and shifted to different extents by altering hyperparameters.

### Policy used for planning

In our work, we assumed that the policy used within the planning loop (i.e. the policy from which actions were sampled during a rollout) was the same as the policy used for sampling actions when actually interacting with the environment. We did this both for simplicity of exposition and computation, and because we think it is likely to be a reasonable approximation to how humans plan. However, there is in theory nothing in our model that prevents the rollout policy from differing from the action policy. In this case, rollouts can still be used to estimate gradients of the future reward with respect to the hidden state, provided that the policy from which rollout actions are sampled is known. This could be done e.g. through the use of importance sampling for off-policy learning. Additionally, in the case of sequential replays, it is plausible that previous replays directly affect future replays, e.g. in a process of exploration. In our current model, there was no option to systematically explore, and previous replays only affected future replays through their effect on the base policy. In theory, it would also be possible to more systematically explore the state space using sequential replays, and indeed we did experiment with ‘rollouts’ corresponding to node expansions of more advanced search algorithms, which can similarly be used to drive improved decision making. More generally, it would also be possible to optimize the rollout policy explicitly for planning by differentiating through the rollout process. This is in contrast to the present work, where the rollout policy was tied to the base policy, and rollouts were treated as part of the ‘environment’. This meant that there was no propagation of gradients to allow for explicit adaptation of the policy to be better for planning.

### Feedback from planning

When performing a rollout, the agent received an additional input on the subsequent timestep consisting of (i) a flattened array of the simulated actions, and (ii) a binary input indicating whether or not the rollout reached the (imagined) goal. Another reasonable choice of feedback input would be to return the sequence of *states* instead of the sequence of *actions*, or potentially to return both. Our reason for favoring the action sequence was primarily that the action space is lower dimensional (4) than the state space (16), which means that the input dimensionality is much lower than it would have been for the state sequence, assuming a one-hot encoding. This does raise the question of where this action sequence would emerge in biological circuits, given that hippocampal replays are canonically assumed to contain spatial information. However, we consider it reasonable that this state information could be converted to information about the actions that would take you there. Instead of returning a binary input of whether the goal was reached, the rollout process could also return the output of a learned value function. We did experiment with returning both the binary ‘goal’ feedback and the imagined value function, and we found that the agent predominantly used the goal information in this case. We therefore chose to remove the learned value from the feedback to simplify the model. This choice is also consistent with previous work in the psychology literature suggesting that human decision making relies on binary evaluations of the success of mental samples (Stewart et al., 2006). However, we imagine that returning a learned value function would be useful in more complicated tasks with multiple or non-binary rewards.

### Stochastic environments and multiple goals

For simplicity, we assumed that the environment was deterministic and that there was only a single goal. However, our model could also be extended to the setting of stochastic environments and multiple or non-binary rewards. In the case of stochastic environments, the agent would still need to simulate a *sample* from the policy. The internal world model was already trained to generate a distribution over new states, and in stochastic environments, we would want to sample from this distribution instead of using the maximum likelihood next state. Provided that the agent has learned a well-calibrated distribution over state transitions, the resulting rollout should still provide an unbiased estimate of the gradient of expected future reward with respect to the hidden state. In the case of multiple goals, it would still be possible to use the agent as-is and return a binary indicator of whether the agent reached any (or each) goal. However, as noted above, it would also be possible to return a learned value estimate instead of the binary goal information. In cases where these goals do not lead to random teleportation, it could also be useful to let the rollout continue beyond the goal. We chose not to do so in the present work, since the transition after reaching the goal was entirely unpredictable, so the simulated action sequence beyond this point would not be informative of expected reward.

### Space in which to plan

We chose rollouts to occur in the space of states and observations. More specifically, the agent had to predict the upcoming state ***s***_*k*_, and a new observation ***x***_*k*_ was constructed automatically from ***s***_*k*_ during the rollout. An alternative would have been to directly learn to predict ***x***_*k*_, which we decided not to do since the majority of the input was constant within an episode. However, in more general task settings, where the environment is more variable, it might be simpler to predict the input directly. Additionally, in partially observable environments, there is a weaker correspondence between states and observations, and rollouts would require samples from the distribution over possible observations. This could either be done directly in observation space or indirectly via some inferred (or known) latent state space.

We consider it likely that humans do not plan explicitly in pixel space and instead use some form of latent planning representation. In the present work, this was also the case to some extent, since the agent input was already an abstract representation. However, in future work, it could be interesting to use a learned latent space instead. This could e.g. be done by training an autoencoder to reconstruct the state and reward information as in the VariBAD model (Zintgraf et al., 2019). Alternatively, planning could take place in a latent space explicitly optimized to yield good plans as in MuZero (Schrittwieser et al., 2020). We did not experiment with any of these possibilities but believe that the results would be comparable to our present work. A major reason for our choice to implement planning in the space of state transitions is that performing high-fidelity rollouts in state space only requires the agent to learn a state transition function. As has been demonstrated in previous work, a transition function could feasibly be learned in a self-supervised manner (Whittington et al., 2020), allowing agents to learn how to plan with little task-specific information. Additionally, rollouts in state space have close parallels to hippocampal replays as detailed in Figure 4.

### Choice of task

The task used for human behavioral experiments and RL agents differs somewhat from the task used for the hippocampal replay data. Notable differences include (i) the presence of ‘away’ trials in the rodent data instead of the teleportation step in the human data, (ii) the different maze sizes and wall configurations, and (iii) the presence of a forced delay between rewards in the rodent data. A natural question is thus why we did not match the human and RL task to the rodent task, which we could not change since this analysis used previously published data. There were a few major reasons for our decision to use different tasks for the humans and RL agents compared to the rodent experiments. One is that the rodent task was not a reaction time task, meaning that there was a forced delay between consecutive rewards. If we introduced a similar delay in the human task, there would be less incentive to act fast. Unfortunately, since we do not have access to intracortical recordings from the human subjects, the speed of acting is the major signal we analyse from our human participants, and it is therefore necessary with a reaction time task. Of course we could still have included away trials and used similar arenas without enforcing a delay between rewards. However, we cared mostly about the ‘home’ trials and therefore saw no reason to make participants spend half their time on ‘away’ trials. Additionally, the smaller Euclidean maze used in the rodent experiments would likely have been too simple for humans and reduced our signal to noise ratio, since there would be less time spent thinking. A simpler task and arena might similarly be simple enough that our RNNs could solve it in a fully ‘model-free’ manner without relying on rollouts to the same extent as in the present work. It is interesting to note that such suboptimality is a key factor of our results, but we believe that this is representative of human behavior as well, where thinking is mostly utilized in scenarios where we do not already know what to do.

### Regularizing time or energy

In our RL agent, we did not incorporate any explicit energy costs for either actions or rollouts. Instead, the only unit of ‘cost’ was time elapsed. We did this since the only explicit incentive to be efficient in our human task was that fast decision making and good actions left more time for collecting reward. It could of course be argued that there is also some energy cost associated with taking actions in our online task, but (i) this energy cost is likely to be negligible, and (ii) if we wanted to model such energy costs in the RL agent, it would require us to introduce an additional hyperparameter to convert between ‘energy’ and ‘time’. We considered it more interpretable and robust to only operate in the space of time, and we also believe that this is representative of many tasks encountered in our daily lives.

### Biological interpretation of the RNN

As mentioned in the main text, we use ‘prefrontal cortex’ to refer to a broader prefrontal network consisting of PFC itself and associated areas of the basal ganglia and thalamus. This follows the work of Wang et al. (2018), which suggested that this prefrontal network can be well modeled across a wide range of tasks as a recurrent meta-reinforcement learner. However, referring to the recurrent part of our RL agent as ‘PFC’ should be seen as a *hypothesis* rather than an *assumption* – indeed the computational and data analysis results all hold equally well if some of the functionality of our RNN is instead carried out in e.g. the hippocampal formation.

A functional argument for expecting prefrontal cortex to be important in the computations carried out by the RNN is its important role in meta-cognition (Botvinick and Cohen, 2014). There is also an extensive associated literature on the role of frontal cortex in value coding (Rushworth and Behrens, 2008) and model-based behavior (Killcross and Coutureau, 2003). The combination of these functionalities makes PFC a natural candidate region in our computational model for deciding whether to plan or act as well as coordinating existing policies with new information from the planning process itself.

